# PfATP2 is an essential flippase on the *Plasmodium falciparum* surface that influences parasite sensitivity to antiplasmodial compounds

**DOI:** 10.1101/2024.12.23.630022

**Authors:** Deyun Qiu, Eleanor England, Adele M. Lehane

## Abstract

Antimalarials play a crucial role in the fight against malaria. However, resistance of the most virulent malaria parasite, *Plasmodium falciparum*, to front-line antimalarials is spreading. To identify new antimalarials, millions of compounds have been screened for their ability to inhibit the growth of blood-stage *P. falciparum* parasites. To gain insight into the mode of action of novel compounds and the ease by which parasites can acquire resistance to them, many have been tested in ‘*in vitro* evolution experiments’, in which parasites are exposed to the compound for a prolonged period of time. In a recent study, parasite resistance to two compounds, MMV007224 and MMV665852, was associated with amplification of the *pfatp2* gene, implicating PfATP2, a putative phospholipid flippase, as a parasite drug target or resistance determinant. These two compounds, along with MMV665794 (which is structurally related to MMV007224) had previously been reported to dysregulate pH in parasites. Here, we show that PfATP2 localises to the parasite surface and is essential for parasite growth. We demonstrate that parasites genetically engineered to overexpress PfATP2 display reduced sensitivity to MMV665794, MMV007224 and MMV665852 compared to parasites with a normal expression level of the protein, and that parasites in which PfATP2 is knocked down become hypersensitive to the three compounds. We show that PfATP2 expression level does not affect the cytosolic pH of parasites, or the potency by which MMV665794 or MMV007224 dysregulate parasite pH. We show that PfATP2-overexpressing parasites internalise a fluorescent phosphatidylserine analogue (NBD-PS) at a greater rate than parasites with a normal expression level of PfATP2, and that parasites in which PfATP2 is knocked down have a reduced rate of NBD-PS uptake. Further, we provide evidence that MMV665794 and MMV007224 inhibit both ATP-dependent and ATP-independent NBD-PS internalisation mechanisms, the former at lower concentrations. Together, our data are consistent with PfATP2 serving as a major ATP-dependent phosphatidylserine internalisation mechanism at the parasite plasma membrane, and being a target of MMV665794 and MMV007224.

## Introduction

*Plasmodium* parasites caused an estimated 249 million cases of malaria and killed 608,000 people in 2022, with *Plasmodium falciparum* causing the majority of deaths (1). *P. falciparum* parasites are adept at developing resistance to the drugs used against them (2,3), necessitating the continual development of new drugs and investigation of new drug targets. Transporters - proteins or protein complexes (including channels, carriers and pumps) that translocate solutes across membrane bilayers or that move lipids from one membrane leaflet to the other - are overrepresented as drug targets and resistance mediators in *P. falciparum*. In a 2018 study, *in vitro* evolution experiments (wherein parasites were exposed to a compound of interest over a prolonged period of time) were performed with 37 antimalarial lead compounds, and the genomes of 262 parasite clones that had acquired resistance were sequenced (4).

Changes (amplification or mutation) in 35 genes were associated with parasite resistance more than once (4). Among these 35 genes, 17 (i.e. 49%) are known or predicted to encode transporters, despite transporters accounting for only ∼ 2.5% of genes in the *P. falciparum* genome (5). Some of these transporters confer resistance by translocating drugs away from their site of action (e.g. PfMRP1 (6)), while others perform essential functions that are inhibitable by small molecules (i.e. serve as drug targets; e.g. PfATP4 (7,8) and PfFNT (9,10)). It has been suggested that some transporters may double as resistance mediators and drug targets (e.g. PfABCI3 (11) and PfCRT (12,13)).

Among the ∼ 117 transporters in *P. falciparum*, 13 belong to the P-type ATPase superfamily (5), members of which can be divided into different types (14,15). P-type ATPases hydrolyse ATP, form a phosphorylated intermediate during their transport cycle, and typically serve as either cation transporters or lipid flippases (15). One of the *P. falciparum* P-type ATPases, PfATP4 (a Type II P-type ATPase), has emerged as a promising drug target. PfATP4 is believed to extrude Na^+^ ions from the parasite cytosol while importing H^+^ (8). A host of chemically diverse molecules have been reported to target PfATP4, including the clinical candidate cipargamin (7,8,16–23).

A role for the Type IV P-type ATPase (P4-ATPase) PfATP2 in parasite susceptibility to antimalarial candidates was uncovered for the first time by Cowell *et al.* (4). P4-ATPases aid in the maintenance of membrane asymmetry by ‘flipping’ specific phospholipids (such as phosphatidylserine (PS) and phosphatidylethanolamine (PE)) from the exoplasmic leaflet of membranes to the cytoplasmic leaflet (reviewed in (24,25)). Typically, PS and PE are more abundant in the cytoplasmic leaflet of membranes (the inner leaflet in the case of the plasma membrane) as a result of the activity of P4-ATPases, while phosphatidylcholine (PC) is more abundant in the exoplasmic leaflet (26). A recent study provided evidence that the homologue of PfATP2 in the rodent-infecting malaria parasite *Plasmodium chabaudi* (PcATP2), when expressed alongside the ‘β subunit’ PcCDC50B, functions as a phospholipid flippase (27). ATP hydrolysis by PcATP2 was stimulated by PS and PE, suggesting that these phospholipids are PcATP2 substrates (27).

Amplification of *pfatp2* (PF3D7_1219600) was associated with *P. falciparum* resistance to the Medicines for Malaria Venture (MMV) compound MMV665852 and was one of several changes observed in parasites resistant to MMV007224 (4). These two compounds, along with a compound called MMV665794 that is structurally similar to MMV007224, are contained within the MMV’s ‘Malaria Box’ compound collection. In other organisms, changes that reduce the function of P4-ATPases or their β subunits in the plasma membrane have been associated with resistance to phospholipid-like pharmacological agents that are thought to rely on P4-ATPase activity for cellular internalisation (28,29). In the plant species *Arabidopsis thaliana*, P4-ATPases have been reported to confer resistance to certain (non-phospholipid-like) toxins by mediating their transport via vesicles to vacuoles, where they are sequestered and degraded (30). The mechanism by which *pfatp2* amplification reduces parasite susceptibility to MMV665852 and MMV007224 is not yet known. In a genome-wide transposon mutagenesis study, PfATP2 was predicted to be essential for the viability of asexual blood stage parasites (31), raising the possibility that PfATP2 is the target of MMV665852 and MMV007224 (4).

MMV007224, MMV665794 and MMV665852 were found previously by our group to be ‘pH dysregulators’ (10). The compounds gave rise to a decrease in the pH of the (normally slightly alkaline) parasite cytosol and an increase in the pH of the (normally acidic) digestive vacuole (DV) (10). The effects of these compounds on parasite pH are consistent with the compounds having protonophore activity (i.e. increasing the permeability of membranes to H^+^). The three MMV compounds (and various synthetic derivatives of them) have also been shown to possess activity against *Schistosoma* worms (e.g. (32–35)).

Here we generated *P. falciparum* lines with altered *pfatp2* expression levels and developed an assay to study PfATP2 activity, in order to determine the function of PfATP2 and understand the mechanism underlying the influence of PfATP2 on parasite susceptibility to MMV665794, MMV007224 and MMV665852.

## Results

### PfATP2 localises to the parasite surface and is essential for parasite viability

To investigate the localisation of PfATP2 and determine whether its expression is important for parasite growth, we made parasite lines (‘PfATP2-GFPreg’ and ‘PfATP2-HAreg’) in which PfATP2 was C-terminally fused to a GFP tag or an HA tag and a *glmS* ribozyme (**Fig. S1**). The *glmS* ribozyme mediates the degradation of the transcript upon parasite exposure to glucosamine (GlcN) (36). PfATP2-GFP and PfATP2- HA primarily localised to the surface of intraerythrocytic trophozoite-stage parasites, with some internal localisation also evident (**Fig. 1A**). Subsequent experiments focused on PfATP2-HAreg parasites. To determine whether PfATP2-HA localises to the parasite plasma membrane or the (closely apposed) parasitophorous vacuole membrane, we investigated its localisation in schizont-infected erythrocytes. The parasitophorous vacuole membrane (marked by EXP2 in **Fig. 1A**) surrounds the developing daughter parasites, whereas PfATP2-HA appears to be on the plasma membrane surrounding each of the daughter parasites (**Fig. 1A**). Upon exposure of PfATP2-HAreg parasites to GlcN (5 mM), we observed a progressive reduction in *pfatp2* mRNA (**Fig. 1B**, red) and loss of parasite viability (**Fig. 1B**, black). Western blotting revealed a reduction in PfATP2-HA protein in PfATP2-HAreg parasites exposed to GlcN (**Fig. 1C**). Thus, our data indicate that PfATP2 is an essential protein localised primarily to the parasite plasma membrane.

**Fig. 1.**
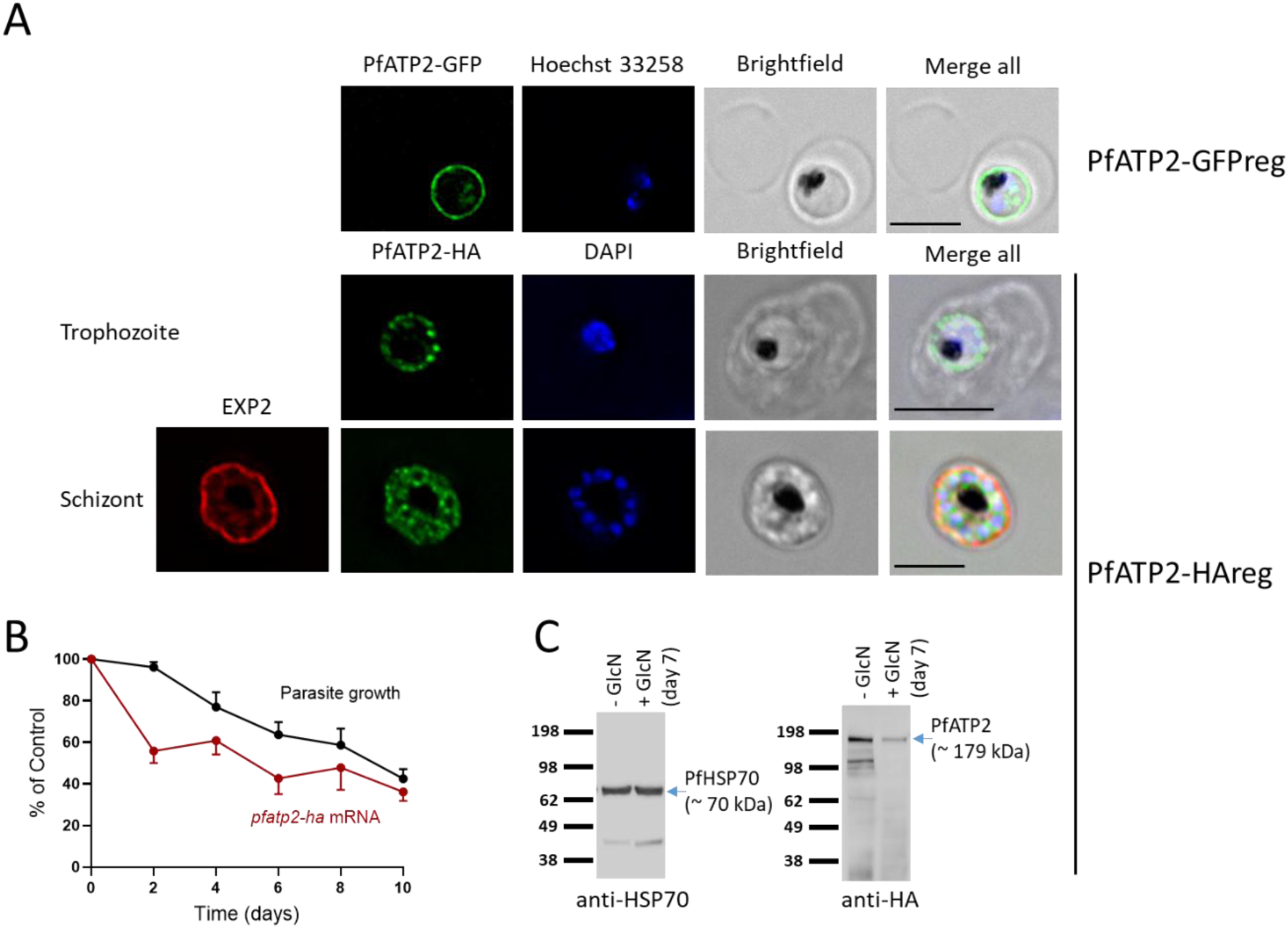
Localisation and knockdown of PfATP2. **A.** PfATP2-GFP (top) and PfATP2-HA (middle) localise to the parasite surface in intraerythrocytic trophozoite-stage PfATP2-GFPreg and PfATP2-HAreg parasites. In segmenting PfATP2-HAreg parasites (bottom), PfATP2-HA appears to localise to the plasma membrane bounding daughter parasites. Scale bars = 5 μm. **B.** GlcN (5 mM) exposure reduces the level of pfatp2-ha mRNA in PfATP2-HAreg parasites (red; with expression compared to the average of two internal controls: 18S rRNA and stRNA) and parasite growth (black; as determined by measuring the parasitaemia at the different time points using flow cytometry). The data are from four independent experiments (except for the mRNA data for Days 2, 4 and 8, which are from three independent experiments). In both cases, the data are expressed as a percentage of those obtained for the same parasites that were not treated with GlcN. **C.** Western blot using antibodies against PfHsp70 (loading control; left) and the HA tag fused to PfATP2 (right) revealed a reduction in PfATP2-HA expression in PfATP2-HAreg parasites treated with 5 mM GlcN for seven days.

### PfATP2 expression level affects parasite sensitivity to MMV665794, MMV007224 and MMV665852

*pfatp2* amplification has been associated with reduced parasite susceptibility to MMV007224 and MMV665852 (4). We used genetically modified lines to directly test whether *pfatp2* expression level affects parasite susceptibility to these compounds and MMV665794. First we generated parasites in which an additional copy of the *pfatp2* gene (untagged) is expressed episomally under the control of the *pfcrt* promoter (‘3D7-PfATP2+’). Relative to control parasites transfected with the empty vector (‘3D7-EV’), 3D7- PfATP2+ parasites (at the trophozoite stage) produce 1.5 ± 0.1 fold more *pfatp2* mRNA (mean ± SEM; n = 5). 3D7-PfATP2+ parasites were slightly less susceptible to MMV665794, MMV007224 and MMV665852 (but not to chloroquine) compared to 3D7-EV parasites (**Fig. 2**), with the IC_50_ values for the three MMV compounds for 3D7-PfATP2+ parasites being 1.4-1.5 fold higher than those for 3D7-EV parasites (**Table 1**).

**Fig. 2:**
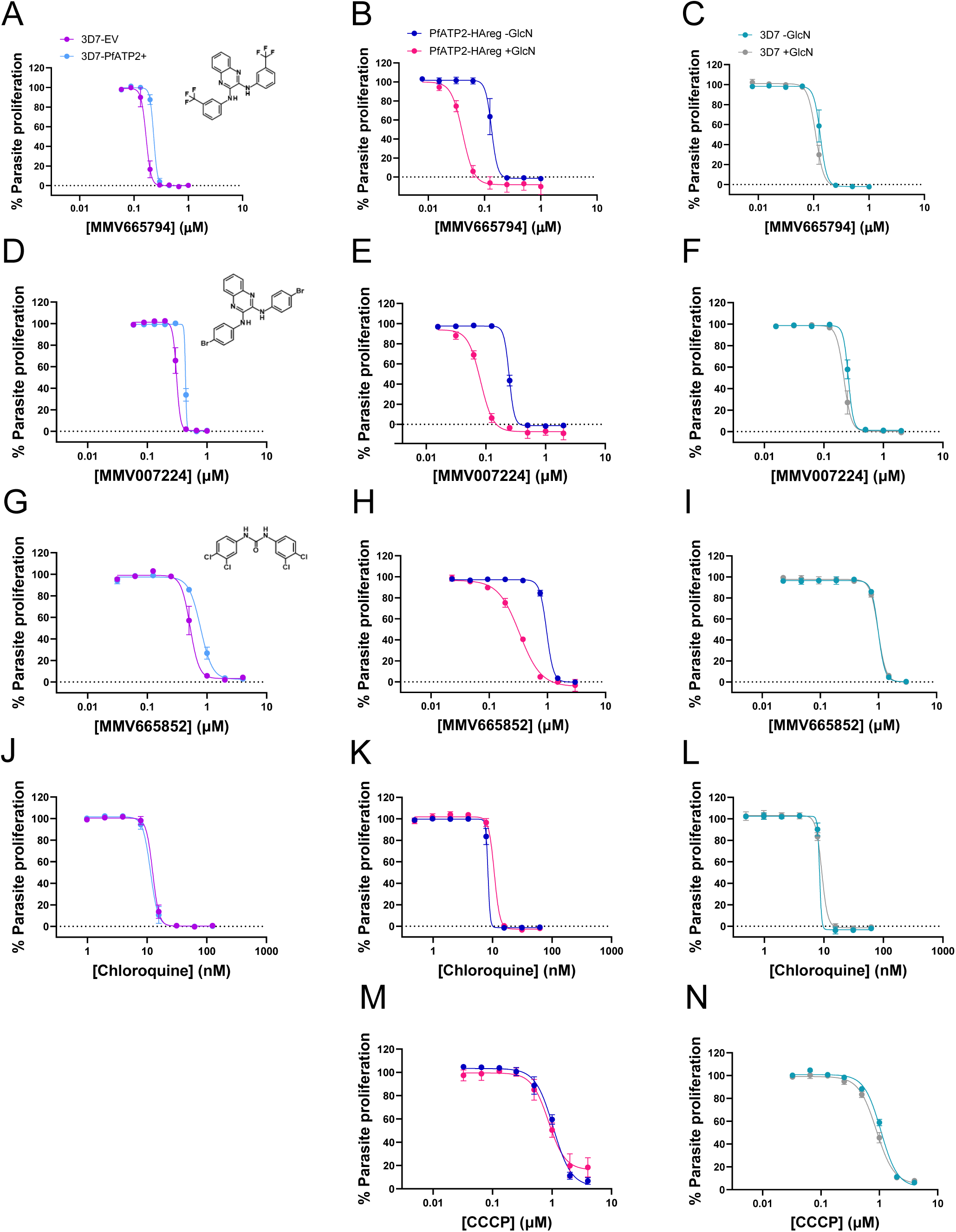
Effect of PfATP2 expression level on parasite response to MMV665794 (A-C), MMV007224 (D-F), MMV665852 (G-I), chloroquine (J-L) and CCCP (M,N). Inhibition of parasite proliferation by the compounds is shown for pfatp2-overexpressing parasites (3D7-PfATP2+; light blue) and empty vector control parasites (3D7-EV; purple) (panels **A,D,G,J**; CCCP was not tested); PfATP2-HAreg parasites that were not exposed to GlcN (-GlcN; blue) or in which PfATP2-HA was knocked down (PfATP2-HAreg +GlcN; pink) (panels **B,E,H,K,M**); and 3D7 parasites that were either exposed (grey) or not exposed (teal) to GlcN (panels **C,F,I,L,N**). The data shown are the mean ± SEM from the number of independent experiments indicated in **Table 1**. Where not shown, error bars fall within the symbols. Where present, GlcN (5 mM) was added to cultures four days before the start of the experiment and maintained throughout the 72 h assay.

**Table 1:** IC_50_ values for growth inhibition of parasites with varying pfatp2 expression levels by the compounds for which dose-response curves are shown in. **Figure 2**. The IC_50_ values in nM (mean ± SEM, from the number of independent experiments stated in brackets) are shown. The fold change (FC) values were calculated by dividing the IC_50_ values obtained for 3D7-PfATP2+, PfATP2-HAreg +GlcN and 3D7 +GlcN parasites by those for 3D7-EV, PfATP2-HAreg-GlcN and 3D7-GlcN parasites, respectively. A FC > 1 indicates resistance and a FC < 1 indicates hypersensitivity. Two-way ANOVAs with Bonferroni’s multiple comparisons tests were performed using the natural logarithm transformed IC_50_s. ns, P ≥ 0.05, *P < 0.05, **P < 0.01, ***P < 0.001. ND, not determined.

| | $IC_{50}$ in nM<br>(mean ± SEM)<br>(n) | | FC (mean<br>± SEM) | $IC_{50}$ in nM<br>(mean ± SEM) (n) | | FC | $IC_{50}$ in nM<br>(mean ± SEM)<br>(n) | | FC |
| --- | --- | --- | --- | --- | --- | --- | --- | --- | --- |
|  | 3D7-EV | 3D7-<br>PfATP2<br>+ |  | PfATP2-HAreg<br>-GlcN | PfATP2-HAreg<br>+GlcN |  | 3D7<br>-GlcN | 3D7<br>+GlcN |  |
| MMV665794 | 172 ±<br>13 (5) | 233 ±<br>14 (5) | 1.37 ±<br>0.10*** | 132 ± 13<br>(4) | 41.4 ±<br>1.0 (4) | 0.323 ±<br>0.030<br>*** | 132 ±<br>11 (4) | 111 ±<br>9 (4) | 0.84 ±<br>0.03<sup>ns</sup> |
| MMV007224 | 312 ±<br>14 (3) | 428 ±<br>11 (3) | 1.38 ±<br>0.09** | 250 ± 4<br>(3) | 81 ± 1<br>(3) | 0.324 ±<br>0.004<br>*** | 262 ±<br>11 (3) | 224 ±<br>17 (3) | 0.85 ±<br>0.05<sup>ns</sup> |
| MMV665852 | 532 ±<br>57 (3) | 786 ±<br>40 (3) | 1.50 ±<br>0.14*** | 971 ± 42<br>(3) | 341 ± 16<br>(3) | 0.351 ±<br>0.007<br>*** | 991 ±<br>41 (3) | 993 ±<br>44 (3) | 1.00 ±<br>0.01<sup>ns</sup> |
| Chloroquine | 13.4 ±<br>0.8<br>(9) | 12.1 ±<br>0.9 (9) | 0.91 ±<br>0.04<sup>ns</sup> | 9.4 ± 0.5<br>(5) | 11.2 ±<br>0.8 (5) | 1.20 ±<br>0.07* | 10.0 ±<br>0.9 (5) | 9.1 ±<br>0.4 (5) | 0.93 ±<br>0.07<sup>ns</sup> |
| CCCP | ND | ND | ND | 1040 ±<br>17 (3) | 854 ± 67<br>(3) | 0.82 ±<br>0.05<sup>ns</sup> | 1063 ±<br>28 (3) | 892 ±<br>42 (3) | 0.84 ±<br>0.03<sup>ns</sup> |

We also tested the sensitivity of PfATP2-HAreg parasites in which PfATP2-HA was knocked down to the three MMV compounds, as well as to chloroquine and the well-characterised protonophore CCCP (**Fig. 2**, **Table 1**). We performed parasite proliferation assays during the window of time in which the majority of GlcN-treated PfATP2-HAreg parasites remained viable despite having a reduced expression of PfATP2-HA (Days 5-7 of GlcN treatment). PfATP2-HA knockdown parasites became hypersensitive to MMV665794, MMV007224 and MMV665852, with the IC_50_ values for the three compounds for PfATP2-HAreg parasites exposed to GlcN being 0.32-0.35 fold those of Control parasites (PfATP2-HAreg parasites not exposed to GlcN). PfATP2-HAreg parasites exposed to GlcN had a slightly higher IC_50_ for chloroquine than those not exposed to GlcN. There was no difference in their sensitivity to CCCP.

To investigate whether the hypersensitivity to the MMV compounds observed with PfATP2-HAreg parasites in the presence of GlcN was a consequence of PfATP2-HA knockdown, and not an off-target effect of GlcN, we also performed parasite proliferation assays with wild-type 3D7 parasites. Exposure of 3D7 parasites to GlcN did not affect their susceptibility to the MMV compounds or to chloroquine or CCCP (**Fig. 2**, **Table 1**).

We next investigated the speed by which the MMV compounds kill PfATP2-HA knockdown and Control parasites. Consistent with previous findings (37,38), we found that the MMV compounds all affected parasite growth in the first cycle of exposure. The compounds gave rise to similar IC_50_ values when they were washed off after 24 h as they did when present throughout the 72 h assays (**Fig. S2**). The level of hypersensitivity to the compounds observed in parasites in which PfATP2-HA was knocked down was similar regardless of whether or not the compounds were removed after 24 h. After a 24 h exposure, the IC_50_ values for the three compounds for PfATP2-HA knockdown parasites were 0.40-0.43 fold those of Control parasites (**Fig. S2**).

Having established that *pfatp2* expression level affects parasite response to MMV665794, MMV007224 and MMV665852, we proceeded to investigate the mechanism involved. Our mechanistic studies focused on MMV665794 and MMV007224, due to concerns about the solubility of MMV665852 at the concentrations required for the experiments. MMV665852 and various derivatives of it have been reported to have low aqueous solubility (< 1.6 μg/mL (< 4.6 μM) (39)).

### PfATP2 expression level does not affect pH dysregulation by MMV665794 or MMV007224

MMV665794, MMV007224 and MMV665852 have all been shown to affect the pH in the parasite cytosol (pH_cyt_) and DV (10). pH dysregulation is expected to be detrimental to parasite growth (40), raising the possibility that the three MMV compounds kill parasites via this mechanism. Given that *pfatp2* expression level alters the antiplasmodial potency of the compounds, we reasoned that if pH dysregulation were the primary mechanism of action of the compounds, this process should be impacted by the *pfatp2* expression level. We therefore investigated whether PfATP2 expression level affects parasite pH regulation or the potency by which the MMV compounds dysregulate pH_cyt_.

The first series of experiments was performed with isolated trophozoite-stage PfATP2-HAreg parasites that were loaded with the pH-sensitive fluorescent dye BCECF and suspended at 37℃ in pH 7.1 Physiological Saline. Control parasites (-GlcN) and PfATP2-HA knockdown parasites (exposed to 5 mM GlcN for four days in the lead up to the experiment) both maintained a similar, slightly alkaline pH_cyt_ when suspended in pH 7.1 Physiological Saline (**Fig. 3A-C**). For Control parasites the pH_cyt_ was 7.41 ± 0.02 and for PfATP2-HA knockdown parasites the pH_cyt_ was 7.42 ± 0.03 (mean ± SEM, n = 6; *P* = 0.6, paired t-test). Upon inhibition of the V-type H^+^ ATPase, which serves as the parasite’s primary regulator of pH_cyt_ (41), with concanamycin A (100 nM) there was a decrease in pH_cyt_ in Control and PfATP2-HA knockdown parasites, as expected from previous studies with different *P. falciparum* strains (e.g. (22,40)). The well-characterised protonophore CCCP (2.5 μM) also gave rise to a rapid decrease in pH_cyt_ in Control and PfATP2-HA knockdown parasites (**Fig. 3A,B**), consistent with previous studies performed with other *P. falciparum* strains (42).

**Fig. 3:**
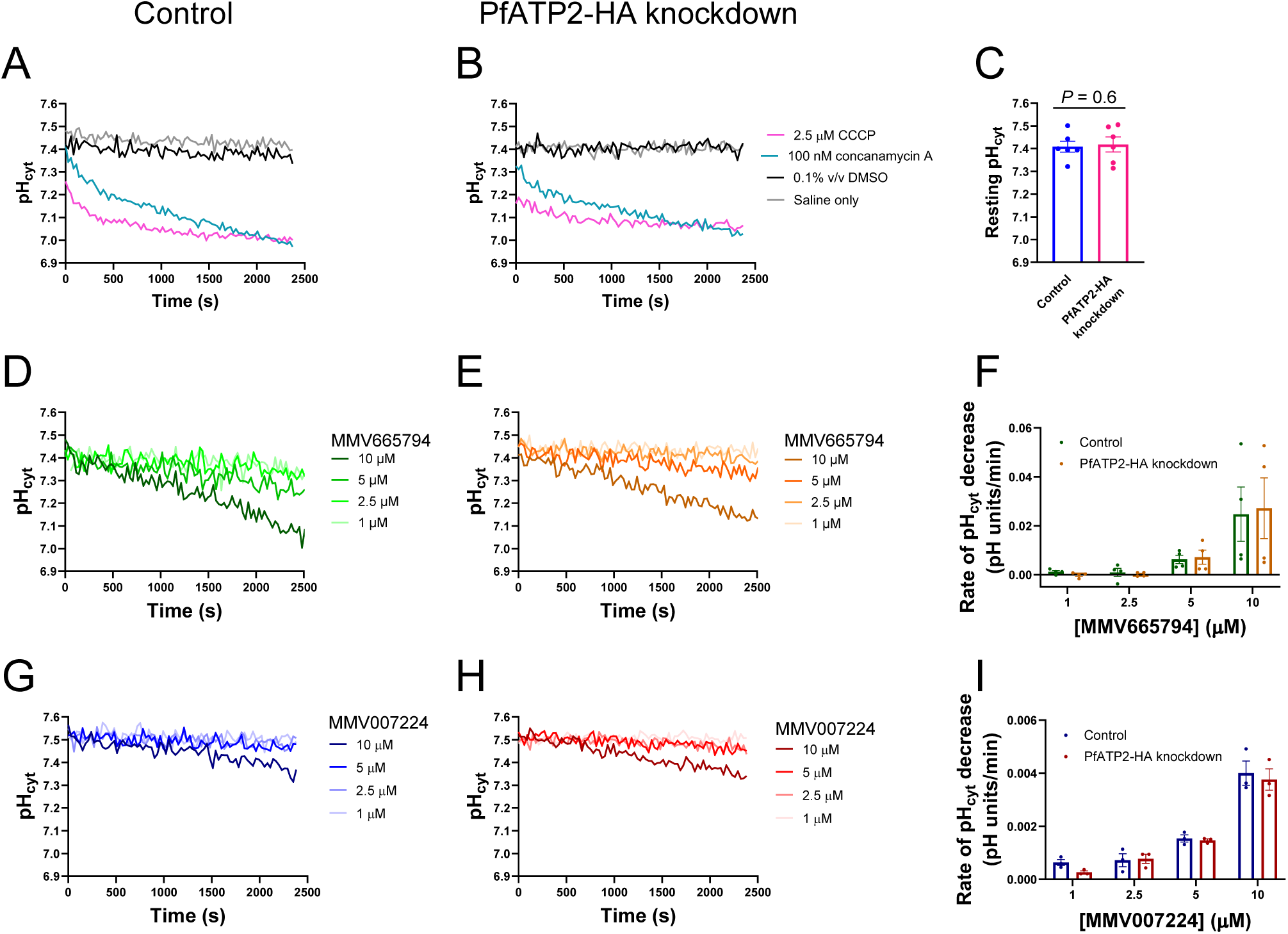
A reduced expression level of PfATP2-HA does not affect the resting pH_cyt_ in P. falciparum parasites or pH_cyt_ dysregulation by MMV665794 or MMV007224. Isolated trophozoite-stage parasites (Control parasites (-GlcN) or PfATP2-HA knockdown parasites (from cultures exposed to 5 mM GlcN for four days) suspended in pH 7.1 Physiological Saline at 37℃ were exposed to either the compound indicated (at the concentration indicated), solvent alone (black; 0.1% v/v DMSO) or Physiological Saline only (grey). GlcN was not present during the measurements. (**A,B,D,E,G,H**) Traces from a single experiment with Control parasites (**A,D,G**) or PfATP2-HA knockdown parasites (**B,E,H**), representative of at least three independent experiments. (C) The resting pH_cyt_ of Control parasites (blue) and PfATP2-HA knockdown parasites (pink) averaged from the first 2000 s of solvent control (0.1% v/v DMSO) traces. The data are from six independent experiments. The resting pH_cyt_ of Control and PfATP2-HA knockdown parasites were compared using a paired t-test, yielding a P value of 0.6. (**F,I**) The rate of pH_cyt_ decrease was estimated from the initial linear portions of the traces for each concentration of MMV665794 (**F**; first 300-2400 s; n = 4) and MMV007224 (I; first 2400 s; n = 3). (**C,F,I**) Control parasites and PfATP2-HA knockdown parasites were tested concurrently in each experiment. The symbols show the data from individual experiments; the bars and error bars show the mean ± SEM. Note that the y-axis ranges are different in panels F and I. Two-way ANOVAs performed with the data shown in Panels F and I revealed a significant effect of the concentration of the MMV compound on the rate of pH_cyt_ decrease (P ≤ 0.04) but no significant effect of PfATP2-HA knockdown (P ≥ 0.5).

We next investigated the effects of MMV665794 and MMV007224 on pH_cyt_ in Control and PfATP2-HA knockdown parasites. In both parasite types, a decrease in pH_cyt_ was readily detectable at concentrations ≥ 5 μM for MMV665794 (**Fig. 3D,E**) and 10 μM for MMV007224 (**Fig. 3G,H**). We estimated the rate of pH_cyt_ decrease from the initial linear portions of the traces. While there was some variability between experiments, the rate of pH_cyt_ decrease was observed to increase with increasing concentrations of the compounds (**Fig. 3F** and **I**). There was no significant effect of PfATP2-HA knockdown on the rate by which the pH_cyt_ decreased in the presence of the compounds.

### MMV665794 and MMV007224 act like protonophores

Protonophores and V-type H^+^ ATPase inhibitors both give rise to a decrease in pH_cyt_ and an increase in DV pH (pH_DV_) (the same effects observed for MMV665794, MMV007224 and MMV665852 (10)). The V-type H^+^ ATPase localises to the parasite plasma membrane, transporting H^+^ out of the parasite, and to the DV membrane, pumping H^+^ into this organelle (41,43,44). The next series of experiments was designed to test whether MMV665794 and MMV007224 have protonophore activity (i.e. render membranes more permeable to H^+^), and if so, to determine whether PfATP2 expression level (in the lead up to the experiment) affected the properties of the plasma membrane in such a way as to influence their potencies as protonophores.

In these experiments, parasites were depleted of ATP (by incubation in Glucose-free Saline) and suspended at time zero in (glucose-free) Low Cl^-^ Saline. The parasite has an acid-loading Cl^-^ transport process, which, under physiological conditions, imports Cl^-^ ions and H^+^ equivalents (45). However, when parasites are suspended in a low Cl^-^ saline (creating an outward Cl^-^ gradient across the parasite plasma membrane), the direction of transport is reversed such that Cl^-^ and H^+^ exit the parasite. Under these conditions, protonophores such as CCCP give rise to an increase in the rate of cytosolic alkalinisation, whereas V-type H^+^ ATPase inhibitors do not (46). ATP-hydrolysing transporters including the V-type H^+^ ATPase and PfATP2 are not expected to be active in the ATP-depleted parasites used in these assays.

In both Control parasites (**Fig. 4A**) and PfATP2-HA knockdown parasites (**Fig. 4B**), CCCP (2.5 μM) increased the rate of cytosolic alkalinisation following suspension of the parasites in Low Cl^-^ Saline. The V-type H^+^ ATPase inhibitor concanamycin A (100 nM) did not affect the rate of cytosolic alkalinisation (**Fig. 4A,B**) – its traces overlapped with those for the solvent control (0.1% v/v DMSO) and Saline only control. This is consistent with the V-type H^+^ ATPase being inactive under the conditions of the experiment.

**Fig. 4:**
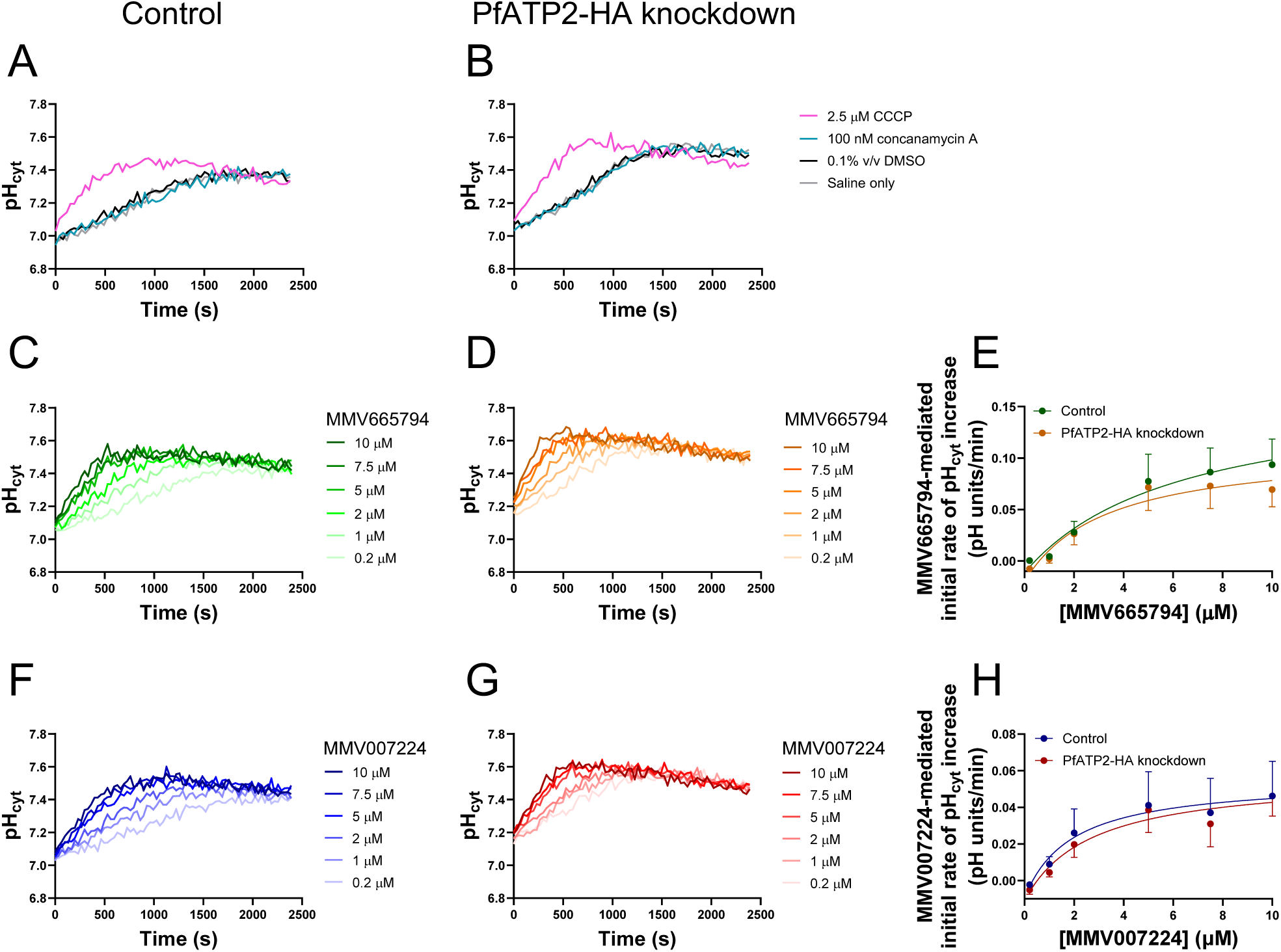
MMV665794, MMV007224 and the protonophore CCCP, but not the V-type H^+^ ATPase inhibitor concanamycin A, increased the rate of cytosolic alkalinisation observed upon creation of an outward Cl^-^ gradient in Control parasites and PfATP2-HA knockdown parasites. (**A-D,F,G**) Representative traces from single experiments with Control parasites (PfATP2-HAreg-GlcN; **A,C,F**) and PfATP2-HA knockdown parasites (PfATP2-HAreg exposed to 5 mM GlcN for four days; **B,D,G**), suspended at time 0 in Low Cl^-^ Saline (37℃). GlcN was not present during the measurements. (**A,B**) Parasites were exposed to either the protonophore CCCP (pink; 2.5 μM), the V-type H^+^ ATPase inhibitor concanamycin A (teal; 100 nM), solvent alone (black; 0.1% v/v DMSO) or Low Cl^-^ Saline only (grey). Each of the four conditions were included in at least two independent experiments, yielding similar results. (**C,D,F,G**) Parasites were exposed to MMV665794 (**C,D**) or MMV007224 (**F,G**), with each compound tested at concentrations of 0.2, 1, 2, 5, 7.5 and 10 μM. The concentration of solvent (DMSO) was 0.1% v/v in each case. (**E,H**) The rate of pH_cyt_ increase was estimated from the initial linear portions of the traces for each concentration of MMV665794 (E; first 150-550 s) and MMV007224 (**H**; first 200-1000 s). Within each experiment, the rate of pH_cyt_ increase calculated for the 0.1% v/v DMSO control trace (first 800-1100 s) was subtracted from the rate of pH_cyt_ increase calculated for each concentration of each compound. Control parasites and PfATP2-HA knockdown parasites were tested concurrently in each experiment. The data are the mean ± SEM from 5 independent experiments. Note that the y-axis ranges are different in panels **E** and **H**. Two-way ANOVAs performed with the data shown in Panels **E** and **H** revealed a significant effect of the concentration of the MMV compound on the initial rate of pH_cyt_ increase (P ≤ 0.0004) but no effect of PfATP2-HA knockdown (P ≥ 0.3).

We next investigated the effects of MMV665794 and MMV007224 on the rate of cytosolic alkalinisation in Control and PfATP2-HA knockdown parasites. Both compounds caused a dose-dependent increase in the rate of cytosolic alkalinisation in Control parasites (**Fig. 4C**,**F**) and PfATP2-HA knockdown parasites (**Fig. 4D**,**G**) suspended in Low Cl^-^ Saline, consistent with the compounds having protonophore activity. For both compounds, the rate of cytosolic alkalinisation increased as the concentration of the compound was increased between 0.2 μM – 5 μM, with the rate of alkalinisation then remaining similar when the concentration was increased further. The initial rate of pH_cyt_ increase mediated by the compounds was higher for MMV665794 (∼ 0.07-0.09 pH units/min at concentrations ≥ 5 μM) than for MMV007224 (∼ 0.04- 0.05 pH units/min at concentrations ≥ 5 μM) (**Fig. 4E**,**H**). There was no effect of PfATP2-HA knockdown on the initial rate of pH_cyt_ increase measured in the presence of the compounds.

In summary, we confirmed that MMV665794 and MMV007224 decrease pH_cyt_ (**Fig. 3**) and provided evidence that they do so by acting as protonophores (**Fig. 4**). Reducing the level of PfATP2-HA expression did not affect the parasite’s resting pH_cyt_ or the protonophore activities of the compounds, suggesting that pH dysregulation may not be the primary mode by which the compounds kill parasites.

### PfATP2 expression level affects NBD-PS internalisation

To determine whether MMV665794 and MMV007224 kill parasites by targeting PfATP2, we first needed to develop an assay to study PfATP2 activity. To investigate whether PfATP2 serves as a phospholipid flippase, we measured the internalisation of a fluorescent PS analogue (NBD-PS) into saponin-isolated trophozoite-stage parasites with varying PfATP2 expression levels. We focused on NBD-PS as a previous study reported that NBD-PS internalisation in isolated *P. falciparum* parasites was sensitive to the P-type ATPase inhibitor vanadate (implicating a P-type ATPase in the uptake process), whereas NBD-PE uptake was reported to be vanadate-insensitive (47). Parasites were incubated with NBD-PS for varying lengths of time, with BSA then used to scavenge the NBD-PS that remained outside the cells or in the extracellular leaflet of the parasite plasma membrane.

PfATP2-HA knockdown parasites (exposed to 5 mM GlcN for two days in the lead-up to the experiment) were found to internalise less NBD-PS over time than Control parasites (-GlcN) (**Fig. 5A**, **Fig. S3A**), consistent with PfATP2 contributing to PS internalisation by the parasites. Visualisation of Control parasites after a 9 min exposure to NBD-PS showed that the majority of the fluorescence was at the parasite surface, consistent with parasite plasma membrane processes playing a major role in NBD-PS internalisation (**Fig. 5B**). To confirm that it was PfATP2-HA knockdown, rather than an off-target effect of GlcN exposure, that led to the reduction in NBD-PS internalisation, we also performed measurements with wild-type 3D7 parasites. GlcN exposure had no effect on NBD-PS internalisation by 3D7 parasites (**Fig. 5C**, **Fig. S3B**).

**Fig. 5:**
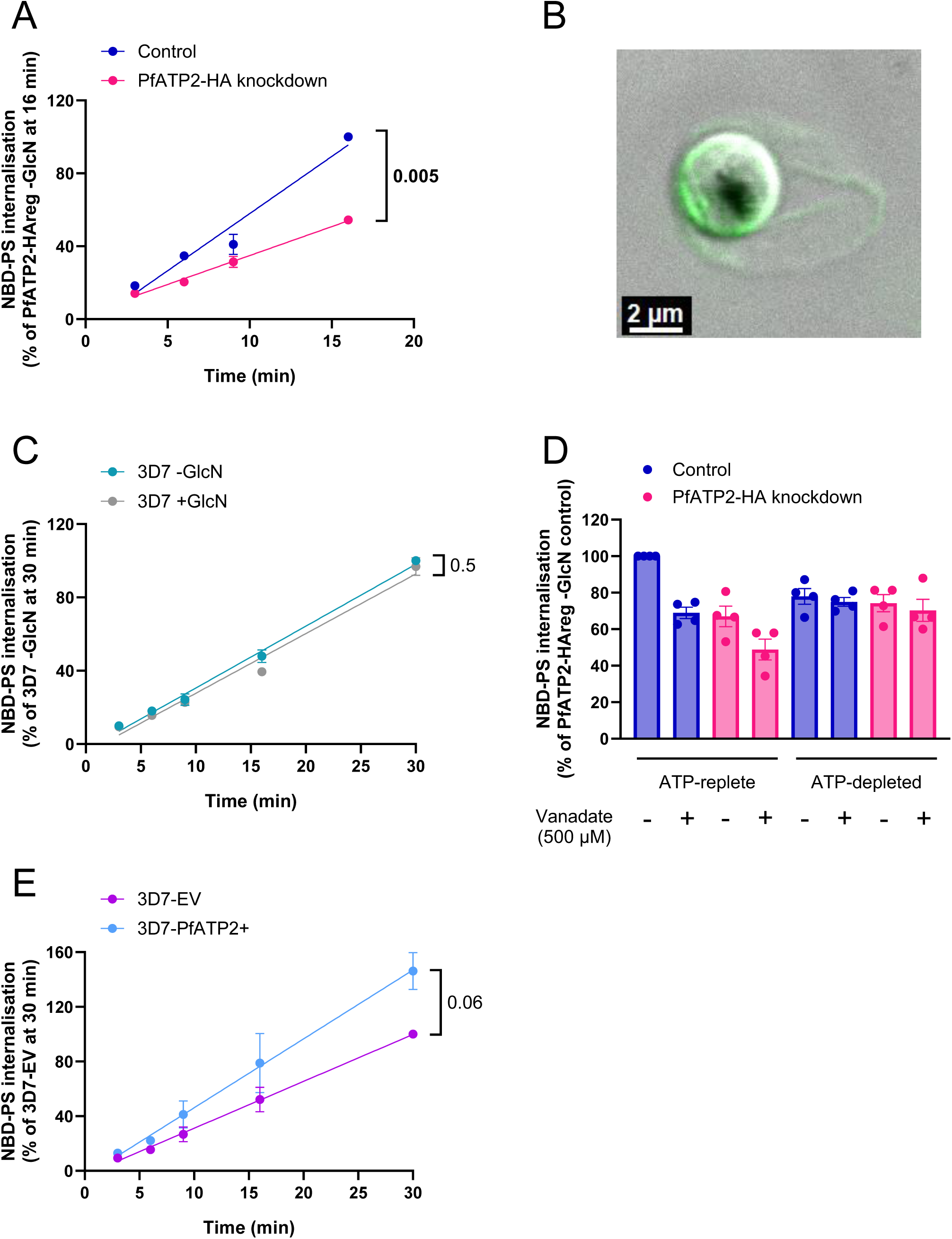
NBD-PS internalisation by parasites is affected by the expression level of PfATP2, the P-type ATPase inhibitor vanadate and the availability of ATP. (**A,C,E**) NBD-PS internalisation over time in PfATP2- HAreg parasites from cultures that were either exposed to 5 mM GlcN for two days in the lead up to the experiment to reduce PfATP2-HA expression (PfATP2-HA knockdown; pink) or that were not exposed to GlcN (Control; blue) (**A**); wild-type 3D7 parasites from cultures that were either exposed to 5 mM GlcN for two days in the lead up to the experiment (+GlcN; grey) or that were not exposed to GlcN (-GlcN; teal) (**C**); pfatp2-overexpressing parasites (3D7-PfATP2+; light blue) and control parasites transfected with the empty vector (3D7-EV; purple) (**E**). The data are the mean (± SEM) from three (**A,E**) or four (**C**) independent experiments. B. Confocal image of an isolated PfATP2-HAreg parasite (-GlcN) after a 9 min incubation with NBD-PS at 15℃. In A-C and E, NBD-PS internalisation was measured in isolated trophozoite-stage parasites suspended in pH 7.1 Physiological Saline at 15℃. **D**. NBD-PS uptake over 9 min at 15℃ in PfATP2-HAreg parasites from cultures that were either exposed to 5 mM GlcN for two days in the lead up to the experiment to reduce PfATP2-HA expression (PfATP2-HA knockdown; pink) or that were not exposed to GlcN (Control; blue), in the absence (solvent control) or presence of vanadate (500 μM). Parasites were either suspended in pH 7.1 Physiological Saline (‘ATP-replete’) or Glucose-free Saline (‘ATP-depleted’). GlcN was not present when parasites were exposed to NBD-PS. The symbols show the data from individual experiments; the bars and error bars show the mean ± SEM. The data are from four independent experiments, in which all the conditions/parasite lines shown were tested concurrently. In **A, C** and **E**, the results of ratio paired t-tests, performed with the slopes obtained when lines were fit to the pre-normalised data (NBD-PS fluorescence (geometric mean)), are shown. For panel **D**, a three-way ANOVA was performed on the natural logarithm transformed pre-normalised data. The effects of vanadate and PfATP2-HA knockdown were both dependent on the presence of ATP (P = 0.02 and P = 0.04, respectively). The pairwise comparisons for which P values are provided in the text are from a post hoc Tukey test. In all panels except **B**, the data are expressed as a percentage of the NBD-PS fluorescence (geometric mean) measured in the parasites/condition indicated on the y axis.

Unless stated otherwise, we used a two day (∼ 48 h) exposure to 5 mM GlcN in the lead up to our NBD-PS uptake experiments (shorter than the exposure times used in the experiments described in the previous sections) to minimise the difference in parasite stage between the PfATP2-HA knockdown and Control parasites. However, using a Coulter Counter, we found that (saponin-isolated) PfATP2-HAreg parasites from cultures that had been exposed to GlcN for two days had a cell volume that was reduced to 79 ± 6% (mean ± SEM, n = 5) of that measured for PfATP2-HAreg parasites not exposed to GlcN (**Fig. S4A**). To investigate whether the smaller volume of PfATP2-HA knockdown parasites might account for their reduced uptake of NBD-PS relative to Control parasites, we also performed experiments in which PfATP2- HAreg parasites were only exposed to GlcN for one day (∼ 24 h). The volume of isolated PfATP2-HAreg parasites from cultures that had been exposed to GlcN for one day was 92 ± 1% (mean ± SEM, n = 3) of that observed for PfATP2-HAreg parasites not exposed to GlcN (**Fig. S4A**). However, the reduction in NBD-PS uptake observed in PfATP2-HAreg parasites that were exposed to GlcN for either one or two days was very similar, with parasites internalising 28-30% less NBD-PS (at the 9 min time point) than Control parasites in both cases (**Fig. S4B**). Thus, the reduction in NBD-PS uptake by PfATP2-HA knockdown parasites cannot be explained by a decrease in cell volume.

As an additional control, we tested PfABCI3-HAreg parasites (made previously (48)) that had been treated with GlcN for one or two days. Like PfATP2-HAreg parasites, PfABCI3-HAreg parasites lose viability over time when exposed to GlcN (48). The effect of GlcN on the volume of PfABCI3-HAreg parasites was similar to that observed for PfATP2-HAreg parasites. The volumes of (saponin-isolated) PfABCI3-HAreg parasites measured after one and two day exposures of parasite cultures to GlcN were 94 ± 2% (mean ± SEM, n = 3) and 82 ± 2% (mean ± SEM, n = 4) those of Control parasites, respectively (**Fig. S4C**). However, unlike PfATP2-HAreg parasites exposed to GlcN for one day, PfABCI3-HAreg parasites exposed to GlcN for one day did not internalise less NBD-PS than PfABCI3-HAreg parasites that had not been exposed to GlcN (**Fig. S4D**). However, NBD-PS internalisation by PfABCI3-HAreg parasites that had been exposed to GlcN for two days was 21 ± 2% lower (mean ± SEM, n = 3) than that observed for PfABCI3-HAreg parasites not exposed to GlcN (**Fig. S4H**). Of note, PfABCI3-HAreg parasites exposed to GlcN lose viability more quickly than PfATP2- HAreg parasites exposed to GlcN (with parasite growth after a two day exposure to GlcN reduced to 87 ± 1% of control levels in the former (48) and 96 ± 2% of control levels in the latter (mean ± SEM, n = 4; **Fig. 1B**)). Whether the finding of reduced NBD-PS uptake by PfABCI3-HAreg parasites after a two day (but not a one day) exposure to GlcN is a result of reduced parasite viability or a direct or indirect effect of PfABCI3 on phospholipid transport is not clear.

To gain further insight into the mechanisms responsible for parasite NBD-PS uptake, we investigated NBD-PS internalisation (at the 9 min time point) by PfATP2-HA knockdown and Control parasites in the presence and absence of the P-type ATPase inhibitor vanadate (500 μM), and under conditions in which the parasites were ATP-replete or depleted of ATP (by incubation in Glucose-free Saline). In ATP-replete parasites, the knockdown of PfATP2-HA or presence of vanadate both resulted in a significant reduction in NBD-PS internalisation (to 67-69% of Control levels; **Fig. 5D**, **Fig. S3F**). In ATP-depleted parasites, neither the knockdown of PfATP2-HA nor the presence of vanadate affected NBD-PS internalisation, as expected (**Fig. 5D**).

Vanadate also reduced NBD-PS internalisation in ATP-replete PfATP2-HA knockdown parasites (*P* = 0.006; **Fig. 5D**). This could be a result of the inhibition of PfATP2-HA, as residual expression of the protein is expected in the knockdown parasites (**Fig. 1B**,**C**), and/or other ATP-dependent internalisation mechanism(s). The finding that NBD-PS internalisation was lower in PfATP2-HA knockdown parasites treated with vanadate than in Control parasites treated with vanadate (*P* = 0.04; **Fig. 5D**) also suggests that at 15℃, the vanadate concentration was not sufficient to completely inhibit ATP-dependent NBD-PS internalisation in parasites expressing a normal level of PfATP2-HA (consistent with the experiments for which data are shown in **Fig. S5B**). The finding that NBD-PS internalisation in ATP-depleted parasites was somewhat higher than in ATP-replete parasites in which PfATP2-HA activity was expected to be minimal (e.g. PfATP2-HA knockdown parasites exposed to vanadate (*P* ≤ 0.02; **Fig. 5D**) or MMV compounds (below; **Fig. S9**)) raises the possibility that the activity of ATP-independent internalisation mechanism(s) is greater in ATP-depleted parasites than in ATP-replete parasites.

Consistent with PfATP2 serving as a PS flippase on the parasite plasma membrane, we found that *pfatp2* overexpressing 3D7-PfATP2+ parasites internalised more NBD-PS than 3D7-EV parasites over the course of 30 min at 15℃ (**Fig. 5E, Fig. S3C**).

Unless stated otherwise, our NBD-PS uptake assays were performed at 15℃. However, similar results were obtained when experiments were performed at 37℃. The overall uptake of NBD-PS was higher at 37℃ than at 15℃, but the effect of knocking down PfATP2-HA on the proportion of NBD-PS taken up was similar, with internalisation reduced by 33 ± 9% at the 9 min time point at 37℃ in the PfATP2-HA knockdown parasites compared to the Control parasites (mean ± SEM, n = 6; *P* = 0.009, ratio paired t-test; **Fig. S5**).

A previous study identified PfCDC50C as the interacting partner of PfATP2 but reported no change in NBD-PS internalisation when PfCDC50C was knocked out (49). Patel *et al.* (49) performed their experiments with parasitised erythrocytes, which were incubated with NBD-PS for 30 min at 37℃. We investigated whether PfATP2 activity could be detected in infected erythrocytes under these conditions. We found that knocking down PfATP2-HA led to a significant decrease in overall NBD-PS internalisation by parasitised erythrocytes, with NBD-PS uptake in parasitised erythrocytes that had been exposed to GlcN for two days being 78 ± 4% of that observed for the-GlcN control cells (mean ± SEM, n = 4; **Fig. S6A**). We also analysed NBD-PS uptake by uninfected erythrocytes (which were distinguishable from the parasitised erythrocytes with flow cytometry). Prior exposure to GlcN had no effect on NBD-PS internalisation by uninfected erythrocytes (**Fig. S6B**).

### MMV665794 and MMV007224 inhibit NBD-PS uptake by parasites

To investigate whether MMV665794 and MMV007224 target PfATP2, we tested a range of concentrations of the compounds (after 9 min at 15℃) for their effects on NBD-PS internalisation. In PfATP2-HAreg Control parasites (-GlcN), MMV665794 significantly inhibited NBD-PS internalisation at concentrations ≥ 5 μM (**Fig. 6A**, **Fig. S3D**), and MMV007224 did so at concentrations ≥ 10 μM (**Fig. 6B**, **Fig. S3E**). These compounds also inhibited NBD-PS internalisation in PfATP2-HA knockdown parasites (**Fig. 6A,B**). This could be a result of the MMV compounds inhibiting residual PfATP2-HA and/or an internalisation mechanism other than PfATP2 (investigated further below).

**Fig. 6:**
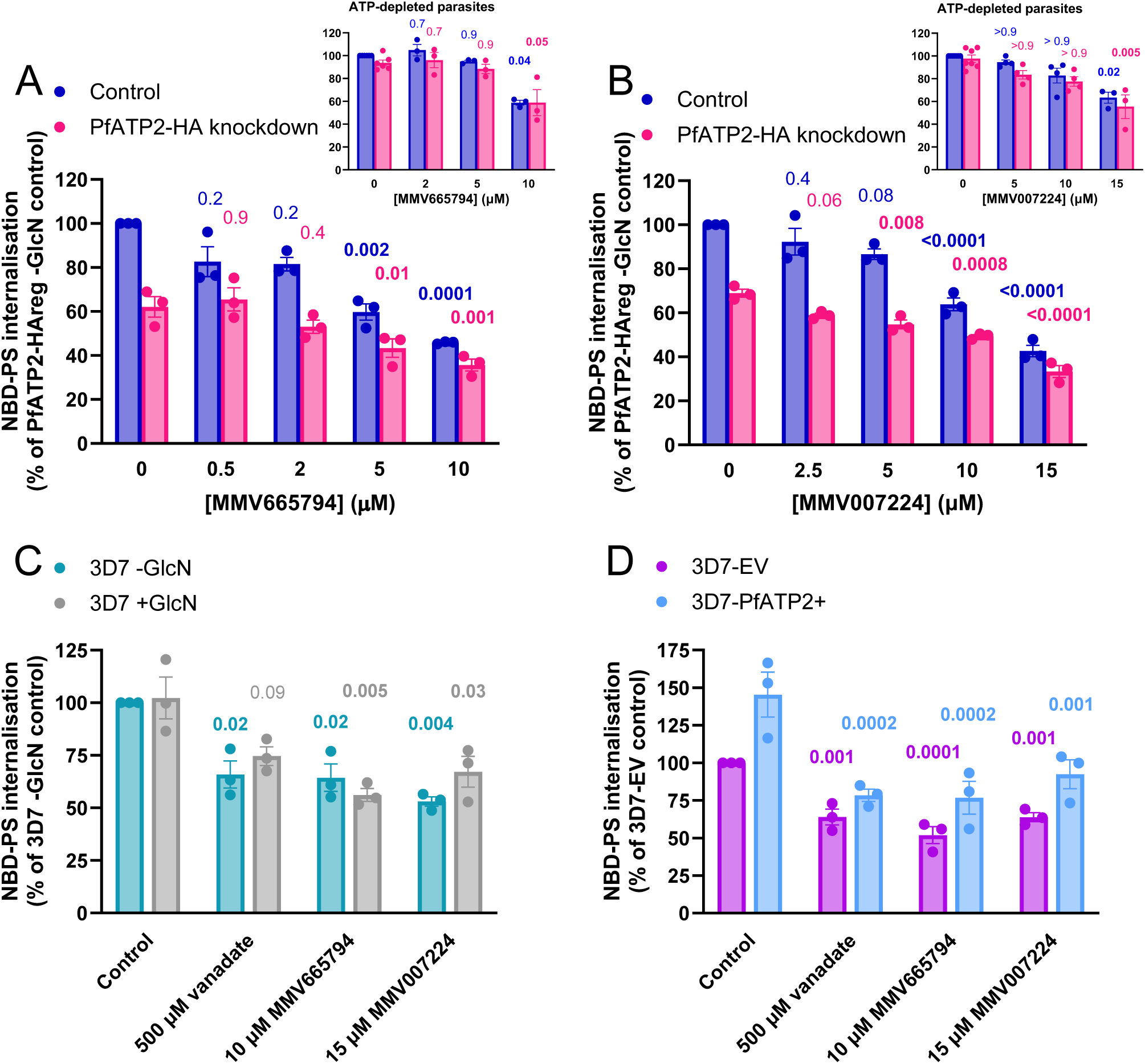
NBD-PS internalisation by parasites is inhibited by MMV665794, MMV007224 and the general P-type ATPase inhibitor vanadate. NBD-PS internalisation was measured in isolated trophozoite-stage parasites suspended in pH 7.1 Physiological Saline. **A** and **B** show data for PfATP2-HAreg parasites from cultures that were either exposed to 5 mM GlcN for two days in the lead up to the experiment to reduce PfATP2-HA expression (PfATP2-HA knockdown; pink) or that were not exposed to GlcN (Control; blue). **Insets:** NBD-PS uptake by ATP-depleted parasites (suspended in Glucose-free Saline). The complete experiments (including data for the ATP-replete parasites tested in parallel) are shown in **Fig. S9**. Panel **C** shows data for wild-type 3D7 parasites from cultures that were either exposed to 5 mM GlcN for two days in the lead up to the experiment (+GlcN; grey) or that were not exposed to GlcN (-GlcN; teal). In **A**-**C**, GlcN was not present when parasites were exposed to NBD-PS. Panel **D** shows data for pfatp2-overexpressing parasites (3D7-PfATP2+; light blue) and control parasites transfected with the empty vector (3D7-EV; purple). In all panels, NBD-PS uptake was measured over 9 min at 15℃ in the absence (solvent control) or presence of the compounds and concentrations indicated. The data are expressed as a percentage of the NBD-PS fluorescence (geometric mean) measured in the parasites/condition indicated on the y axis. The data are from at least three independent experiments, in which all the conditions/parasite lines shown within one panel were tested concurrently (with the exception of the insets). The symbols show the data from individual experiments; the bars and error bars show the mean ± SEM. Statistical comparisons were made on the natural logarithm transformed pre-normalised data (NBD-PS fluorescence (geometric mean)) using two-way ANOVAs with Dunnett’s multiple comparisons tests. The P values for comparisons with the solvent control for the same parasite type are shown, with values indicating statistical significance (P ≤ 0.05) shown in bold.

In wild-type 3D7 parasites that were either exposed to 5 mM for two days prior to the experiment or not exposed to GlcN, MMV665794 (tested at a single concentration of 10 μM), MMV007224 (15 μM), and vanadate (500 μM) all inhibited NBD-PS uptake to a similar degree (to ∼ 60-70% of control levels; **Fig. 6C**).

We also tested MMV665794, MMV007224 and vanadate for their effects on NBD-PS uptake by 3D7- PfATP2+ and 3D7-EV parasites. In the absence of test compound (solvent control), 3D7-PfATP2+ parasites internalised 1.5 ± 0.1 fold more NBD-PS than 3D7-EV parasites at the 9 min time point (mean ± SEM, n = 6; *P* = 0.001, ratio paired t-test; **Fig. 5E** and **Fig. 6D**). All three compounds inhibited NBD-PS internalisation by 3D7-PfATP2+ and 3D7-EV parasites (**Fig. 6D**). In the presence of these inhibitors, NBD-PS internalisation was similar between 3D7-PfATP2+ and 3D7-EV parasites, although still slightly higher in the former, perhaps because the concentrations used were not sufficient to fully inhibit PfATP2 activity in the PfATP2- overexpressing parasites.

While the experiments for which data are shown in **Fig. 6** were performed at 15℃, MMV665794 and MMV007224 also inhibited NBD-PS internalisation at 37℃ (**Fig. S5B**). No difference in MMV665794- or MMV007224-mediated inhibition of NBD-PS internalisation was observed between measurements performed at 15℃ and 37℃ in PfATP2-HAreg parasites (whether or not they had been exposed to GlcN) (**Fig. S5B**). In contrast, vanadate showed greater inhibition of NBD-PS uptake at 37℃ than at 15℃ in Control parasites (but not PfATP2-HA knockdown parasites) (**Fig. S5**). This raises the possibility that vanadate may not be fully soluble at 15℃, and that at this temperature the concentration in solution was not sufficient to fully inhibit P-type ATPase activity in parasites with a normal expression level of PfATP2- HA.

While we used a 10 min pre-incubation time for the MMV compounds (and vanadate) in the experiments for which data are shown in **Fig. 6**, similar results were obtained when the compounds were added to parasites at the same time as NBD-PS (**Fig. S7**).

We tested the protonophore CCCP to investigate whether dysregulation of parasite pH_cyt_ might indirectly affect NBD-PS internalisation. CCCP (tested at 100 nM and 5 μM at 15℃) did not show evidence for the inhibition of NBD-PS internalisation in PfATP2-HA knockdown or Control parasites (**Fig. S8**). Thus, the effects of the MMV compounds on NBD-PS internalisation are unlikely to stem from their effects on pH_cyt_. When tested at the higher concentration, CCCP appeared to increase fluorescence. Whether this resulted from an increase in NBD-PS uptake or a contribution to the fluorescence from CCCP itself is not clear.

We next compared the effects of MMV665794 and MMV007224 on NBD-PS internalisation in ATP-replete and ATP-depleted parasites. If the compounds inhibit NBD-PS uptake solely via inhibition of PfATP2, they should have no effect in ATP-depleted parasites. However, when tested at 10 μM (MMV665794) or 15 μM (MMV007224) (the highest concentrations tested in **Fig. 6**), both compounds reduced NBD-PS internalisation by both ATP-replete and ATP-depleted parasites (**Fig. S9A**, **Fig. 6A**,**B** insets). We next investigated lower concentrations of each compound. When tested at 2 μM and 5 μM (MMV665794; **Fig. S9B**) or 5 and 10 μM (MMV007224; **Fig. S9C**), the compounds did not significantly inhibit NBD-PS internalisation by ATP-depleted parasites (**Fig. 6A**,**B** insets). The data suggest that the compounds inhibit both an ATP-dependent NBD-PS internalisation mechanism (likely PfATP2, with some residual expression expected in the PfATP2-HA knockdown parasites) and an ATP-independent mechanism, and that they inhibit the former more potently.

We also investigated the effects of MMV665794 and MMV007224 on NBD-PS uptake by parasitised erythrocytes and uninfected erythrocytes. NBD-PS internalisation by parasitised erythrocytes was reduced by MMV665794 and MMV007224 (**Fig. S6A**). Both compounds caused a significant reduction in NBD-PS internalisation by uninfected erythrocytes as well (**Fig. S6B**), suggesting that they are also active against one or more PS uptake process(es) in erythrocytes.

Together, the data are consistent with MMV665794 and MMV007224 inhibiting PfATP2, but suggest that they also inhibit other NBD-PS uptake processes at the higher concentrations tested.

## Discussion

This study provides evidence that PfATP2 serves as an essential phospholipid flippase on the *P. falciparum* plasma membrane, with PS being a substrate of the transporter. However, measurements of the activity of the protein in purified form, or expressed in a heterologous expression system, would be required for direct confirmation. PfATP2 has the signature motifs characteristic of a P4-ATPase (50) and contains a Q residue in transmembrane helix 1 equivalent to Q88 in human ATP8A1, which interacts with the headgroup of PS and is critical for PS transport (26,51–54).

Most P4-ATPases interact with a non-catalytic ‘β subunit’ (26). In *P. falciparum* there are three candidate ‘cell division control 50’ (CDC50) proteins. One of these, PfCDC50C, was recently shown to interact with PfATP2 in parasites and to be essential for the viability of blood-stage parasites (49). PfCDC50A was not expressed in asexual parasites and PfCDC50B interacted instead with guanylyl cyclase α (GCα) (49). Studies with the related apicomplexan parasite *Toxoplasma gondii* yielded consistent findings, with the *T. gondii* homologue of PfATP2 (termed ATP2B (55) or TgP4-ATPase1 (50)) found to interact with the homologue of PfCDC50C (termed CDC50.4 (55) or TgLem1 (56)) (55,56). In a study with recombinantly expressed *P. chabaudi* proteins, PcATP2 was found to form a heterodimer with PcCDC50A and PcCDC50B (the expression of PcCDC50C in yeast was low, precluding further investigation) (27). PcATP2/PcCDC50B was reported to hydrolyse ATP in the presence of PS and PE (27). Together, these findings suggest that PfATP2 might be able to form an active complex with more than one of the PfCDC50 proteins *in vitro*, but that within the parasite it primarily associates with PfCDC50C.

It was reported that the depletion of PfCDC50C by conditional gene disruption did not impact the uptake of NBD-PS (or NBD-PE or NBD-PC) by parasitised erythrocytes. In contrast, we saw a significant effect of knocking down PfATP2 on NBD-PS uptake, even when we attempted to replicate the conditions used by Patel et al. (49) (i.e. using parasitised erythrocytes rather than isolated parasites and allowing 30 min at 37℃ for NBD-PS uptake). How these different findings can be reconciled is not yet clear. One possible explanation might relate to differences in parasite stage, which may affect the overall contribution of PfATP2 to NBD-PS uptake by parasitised erythrocytes. Our measurements were performed with erythrocytes infected with mature trophozoites that were larger than those for which images are shown in the study by Patel et al. (49).

It remains to be determined whether PS is the sole substrate of PfATP2 or whether it has additional substrates. PcATP2 was suggested to transport both PS and PE (27), whereas ATP2B was found to serve as a PS flippase in the *T. gondii* plasma membrane (55,56) but was reported not to transport PE (56).

Furthermore, it has been shown that PS internalisation by isolated *P. falciparum* parasites was partially inhibited by the P-type ATPase inhibitor vanadate but that PE internalisation was not (47). In other organisms, a range of substrates have been identified for P4-ATPases including PS, PE, PC, glucosylceramide and galatosylceramide (reviewed in (26)). The *in situ* assay developed to measure PfATP2 activity in this study could be useful to further investigate the substrate range and transport properties of PfATP2. The assay can also be used to study the regulation of PfATP2 – for example to test whether its activity is stimulated by signalling molecules such as phosphatidylinositol-4-phosphate (PI4P) (as observed for PcATP2 (27) and also expected for PfATP2 based on the presence of a GYAFS motif (50,57)).

P4-ATPases and the membrane asymmetry they bring about have been found to contribute to numerous processes in diverse cell types, regulating the biophysical properties of membranes and playing an important role in protein-membrane interactions, cell signalling, vesicle-mediated transport and (in some cases) lipid scavenging from a host (26,50). In *T. gondii*, ATP2B/CDC50.4 was found to have a particularly important role in the secretion of microneme contents (55), motility, parasite egress from host cells and invasion of new host cells, and was not required for normal intracellular growth (56). In contrast, PfCDC50C was found to be important for the maturation of trophozoites within host erythrocytes (49). PfCDC50C was also proposed to play a role in the endocytosis of host haemoglobin (49), suggesting that PfATP2/PfCDC50C-mediated PS asymmetry may be important for this process. *P. falciparum* is able to synthesise PS, PE and PC from precursors and to (directly or indirectly) interconvert between them (58,59); thus, it is not clear whether scavenging PS from the host would be an essential role for PfATP2 in blood-stage parasites.

Our finding that PfATP2-HA knockdown substantially reduces NBD-PS internalisation in ATP-replete (but not ATP-depleted) parasites suggests that PfATP2 is a major contributor to ATP-dependent PS uptake in parasites. However, contributions by other proteins cannot be ruled out. In addition to PfATP2, the *P. falciparum* genome is predicted to encode three other putative P4-ATPases (PfATP7, PfATP8 and PfATP11) and two proteins that contain both P4-ATPase and guanylyl cyclase domains (GCα and GCβ) (5). Only PfATP2, PfATP8 and PfGCα were predicted to be essential in asexual blood-stage parasites (31), with PfGCα subsequently confirmed to be essential (for parasite egress) (60) and PfATP2 now confirmed to be essential in this study. The localisation of PfATP7, PfATP8 and PfATP11 in blood-stage *P. falciparum* parasites has not yet been reported. A study with the murine parasite *P. yoellii* reported that ATP7 is required for parasite survival in the mosquito and that PC is its likely substrate (61). GCα has been reported to be expressed maximally during the schizont stage and to localise to vesicular structures in the cytoplasm (60), while GCβ is dispensible (and not expressed highly) in asexual blood-stage parasites (62).

There remains a significant level of NBD-PS internalisation in ATP-depleted parasites, and in ATP-replete parasites exposed to the P-type ATPase inhibitor vanadate, suggesting that one or more ATP-independent processes also contribute to parasite PS uptake. Scramblases are a type of transporter that catalyse bidirectional lipid transport and do not hydrolyse ATP. Blood-stage *P. falciparum* parasites have been shown to express a phospholipid scramblase (PfPLSCR), however it appears to be localised intracellularly in trophozoite-stage parasites and it is not clear whether there is any protein present on the plasma membrane during this stage (63). The contributor(s) to ATP-independent PS internalisation remain to be determined.

When considered together, our findings with MMV665794 and MMV007224 suggest that PfATP2 is likely their primary target in parasites. We found that parasites in which PfATP2 was knocked down became hypersensitive to growth inhibition by MMV665794 and MMV007224 (and MMV665852), and that parasites in which PfATP2 was overexpressed displayed reduced sensitivity to the compounds. We showed that MMV665794 and MMV007224 inhibit NBD-PS internalisation by ATP-replete parasites at lower concentrations than those needed to inhibit NBD-PS uptake by ATP-depleted parasites. The concentrations required to detect inhibition of NBD-PS uptake in our assays were ∼ 20-fold higher than the IC_50_ values of the compounds in (longer-term) growth assays. Unfortunately, we were unable to determine whether MMV665852 also inhibits NBD-PS uptake at 20-fold its IC_50_, as a result of the higher IC_50_ of this compound and its low aqueous solubility (39).

The amplification of *pfatp2* (4) or its 1.5-fold overexpression (this study) only gave rise to a modest reduction in parasite sensitivity to MMV007224, MMV665794 and MMV665852 (∼ 1.5-fold increase in IC_50_). The low level of resistance may reflect the low level of PfATP2 overexpression. It is also possible that the expression of PfCDC50C could become limiting for PfATP2 activity when PfATP2 is overexpressed. It remains to be determined whether parasites can also acquire resistance to the compounds via point mutations in PfATP2 that affect their binding.

Additional detrimental effects of the compounds in parasite membranes may also limit the extent to which changes in *pfatp2* expression can protect parasites from the MMV compounds. All three MMV compounds are lipophilic (with LogP values between 6 and 6.7 in Pubchem; computed using XLogP3 3.0 (64)) and have been found previously to dysregulate parasite pH (10). In this study we found that MMV665794 and MMV007224 are likely to dysregulate pH as a result of having protonophore activity. We found that the concentrations of the compounds that gave rise to a detectable decrease in pH_cyt_ were similar to those that gave rise to a detectable decrease in NBD-PS internalisation. It is possible that pH dysregulation becomes the primary mode of action of the compounds in parasites that overexpress PfATP2. The finding that the MMV compounds can inhibit ATP-independent NBD-PS uptake suggests that the compounds may have additional targets in parasite membranes that have not yet been discovered.

While parasites with low-level resistance to MMV007224 and MMV665852 were generated in the Cowell *et al.* study and associated with amplification of *pfatp2* (4), multiple attempts to generate resistance to MMV665794 through *in vitro* evolution experiments with wild-type parasites (3D7 and Dd2) were unsuccessful (65). However, parasites with low-level resistance to MMV665794 (2-2.5 fold elevated IC_50_s) were subsequently generated successfully by exposing ‘hypermutator’ parasites with mutations in DNA polymerase δ to the compound (65). Attempts to increase the level of resistance further were not successful (65). Interestingly, parasite lines with low-level resistance to MMV665794 were found to have mutations in a gene encoding a protein of unknown function with four predicted transmembrane domains (PF3D7_1359900; ‘Quinoxaline-resistance protein 1’ (QRP1)) (65). Parasites engineered to express the resistance-associated QRP1 mutations (D1863Y or G1612V) were found to display low level resistance to both MMV665794 and MMV007224 (MMV665852 was not tested) (65). Results of a genome-wide transposon mutagenesis study (31), together with the finding of a frameshift mutation in the gene encoding QRP1 in MMV007224-selected parasites (65), provide strong evidence that QRP1 is not essential in blood-stage parasites, and therefore that the primary mode of action of the MMV compounds is unlikely to be inhibition of QRP1 function. Whether QRP1 has a role in regulating PfATP2 activity, or confers resistance to MMV665794 and MMV007224 via a different mechanism, remains to be determined.

MMV007224, MMV665794 and MMV665852 all display toxicity in some assays (66) and would need to be optimised for further development. Whether the compounds’ toxicity against human cells stems from their protonophore activity, inhibition of one or more of the 14 human P4-ATPases or other phospholipid uptake pathways, or different mechanism(s) is not known. We observed that MMV665794 and MMV007224 inhibited NBD-PS internalisation by uninfected erythrocytes, which suggests that the compounds might have activity against one or more ATP-dependent and/or ATP-independent uptake pathway(s) on the erythrocyte plasma membrane. There is evidence for the presence of three P4-ATPases in human erythrocytes: ATP11A, ATP11B and ATP11C, with ATP11C being the most highly expressed (67–69).

Recently, some quinoxaline-based compounds (similar in structure to MMV007224 and MMV665794) active against *Schistosoma* parasites were found to be more potent against *P. falciparum* than the compounds tested in this study, with *pfatp2* amplification and QRP1 mutations again shown to confer low-level resistance (70). Further studies on whether PfATP2 and its homologues in other parasites can be exploited as new drug targets are eagerly awaited. Screening compounds for their activity against the *pfatp2* knockdown line generated here would provide an efficient starting point for pinpointing additional PfATP2 inhibitors, with measurements of NBD-PS internalisation serving as a secondary assay for any ‘hits’ identified.

## Materials and Methods

### Compounds

MMV007224 was purchased from Vitas-M Laboratory. MMV665794, and additional quantities of MMV007224, were kindly provided as powders by MMV. MMV665852 was purchased from Key Organics.

16:0-06:0 NBD PS (1-palmitoyl-2-{6-[(7-nitro-2-1,3-benzoxadiazol-4-yl)amino]hexanoyl}-sn-glycero-3- phosphoserine (ammonium salt); Aventi Polar Lipids) was purchased from Merck.

### Plasmid constructs

To overexpress PfATP2 (Pf3D7_1219600) in *P. falciparum* parasites, cDNA was prepared from 3D7 parasites, and the full length cDNA encoding PfATP2 was amplified with primers P1 and P2 (Table S1), carrying XhoI and KpnI restriction sites, respectively. A double stop codon was added to the 3’ end of the cDNA. The resulting amplicon was then cloned into the pGlux-1 vector (71), which carries a selection marker gene encoding human DHFR, using In-Fusion HD cloning (TAKARA). The *P. falciparum* chloroquine resistance transporter (*pfcrt*) promoter was used to drive *pfatp2* expression.

To engineer the PfATP2-HAreg line, a 996 bp fragment was amplified from a region of the 3D7 *pfatp2* coding sequence immediately upstream of the stop codon using primers P3 and P4, which harbour BglII and PstI restriction sites, respectively. The fragment was ligated into the BglII and PstI sites of an HA glms vector (72).

To engineer the PfATP2-GFPreg line, the same 996 bp fragment was amplified using primers P3 and P5, which contain BglII and KpnI sites, respectively. The fragment was ligated into the BglII and KpnI sites of pGFP_*glmS* (36).

Before transfection, the production of the desired constructs was confirmed by Sanger sequencing at the Genome Discovery Unit - ACRF Biomolecular Resource Facility, The John Curtin School of Medical Research, Australian National University. The primers used for sequencing are shown in Table S1.

### Parasite culture and transfection

The use of human blood in this study (from anonymous donors) was approved by the Australian National University Human Research Ethics Committee (Protocol numbers 2011/266 and 2017/351).

Blood-stage *P. falciparum* parasites (the 3D7 strain, the PfABCI3-HAreg line (48), and the transgenic parasites made in this study) were maintained within human erythrocytes (typically Group O+, 4% haematocrit) in RPMI 1640 medium containing 25 mM HEPES and GlutaMAX^TM^ (ThermoFisher Scientific Cat. # 72400120), and supplemented with 11 mM (additional) glucose, 200 μM hypoxanthine, 24 μg/mL gentamicin and 6 g/L Albumax II (defined as complete medium in this study). Cultures were kept in a horizontally-rotating shaking incubator at 37°C under a low-oxygen atmosphere (1% O_2_, 3% CO_2_ and 96% N_2_). Parasites were synchronised by sorbitol treatment (73).

Transfections were performed when parasites were predominantly at the young ring stage, with 100 μg of the appropriate circular DNA constructs transfected by electroporation as described previously (74). Transgenic parasites were selected with 5 nM WR99210 (Jacobus Pharmaceuticals; applied 48 h post-transfection) (PfATP2-HAreg, 3D7-PfATP2+ and 3D7-EV) or 5 μM blasticidin (PfATP2-GFPreg) and were cultured every second day until parasites resistant to the selection agents were observed. For PfATP2- HAreg and PfATP2-GFPreg, this was followed by two cycles each consisting of one week with the appropriate selection agent followed by one week without. The presence of integrated transgenic DNA in the PfATP2-HAreg and PfATP2-GFPreg parasites was determined by PCR from total parasite genomic DNA (extracted from isolated trophozoite-stage parasites (see below)) using a QIAGEN DNeasy Plant Kit) using PrimerSTAR GXL DNA polymerase (TAKARA). All primers used for PCRs and sequencing are listed in Table S1. Primer P30 paired with P26 (PfATP2-HAreg) or P27 (PfATP2-GFPreg) were used to investigate whether transfection was successful, and primer P8 paired with P26 (PfATP2-HAreg) or P27 (PfATP2-GFPreg) was used to check for successful integration (with P8/P29 used to detect the unmodified locus). For PfATP2- HAreg and PfATP2-GFPreg, clonal lines were obtained using the FACS-based approach described previously (75). The presence of the desired integration events in the clones were confirmed by PCR using the primers outlined above, and (for PfATP2-HAreg) by sequencing using primer P8.

### Parasite isolation from host erythrocytes

Trophozoite-stage parasites were isolated from their host erythrocytes by adding saponin (final concentration 0.05% w/v) to parasite cultures (∼ 4% haematocrit, 2-10% parasitaemia), inverting the tube several times to mix, then centrifuging immediately (1000 × g, 5 min). The supernatant medium was removed and the parasites were resuspended then washed several times (with 12,000 × g, 30 s centrifugation steps) in ‘bicarbonate-free medium’ (bicarbonate-free RPMI 1640 medium supplemented with 25 mM HEPES, 11 mM (additional) glucose and 0.2 mM hypoxanthine (pH 7.1)) at 37℃ (unless stated otherwise).

### pfatp2 mRNA quantification

Quantitative reverse transcription PCR (qRT-PCR) was performed to quantify the expression level of *pfatp2* mRNA. Total RNA was extracted from isolated trophozoite-stage parasites using the QIAGEN RNeasy mini kit. DNA-free RNA samples (1-2 μg) were reverse-transcribed using oligo dT(18) and random primers (50℃ for 1 h in 20 μL reactions) with SuperScriptIII enzyme as recommended by the manufacturer (Invitrogen). The reaction mixes were then diluted 1:3 in nuclease-free water. qPCR was performed using FastStart Universal SYBR Green Master (ROX) (Roche) in 10 μL reactions, with each reaction containing 5 μL Master mix, 2 μL of diluted cDNA template and 0.5 μM of each oligonucleotide primer (Table S1, P6-P15). The qPCR was performed with the ViiA™ 7 Real-Time PCR System (ThermoFisher Scientific). The 2^-ΔΔCt^ method (76) was used to calculate relative changes in gene expression.

### Monitoring parasite viability by flow cytometry

The effect of PfATP2 knockdown on the survival of PfATP2-HAreg parasites was assessed using a method similar to that described previously (77). On Day 0 (at which time parasites were in the trophozoite stage), parasitemia was accurately determined using a previously described flow cytometry method (75). After determining the starting parasitemia, two 50 mL cultures were set up with PfATP2-HAreg parasites at a parasitemia of 0.5%. GlcN (5 mM) was added to one culture, while an equivalent volume (0.5 mL) of solvent (culture medium without Albumax II) was added to the other. WR99210 (5 nM) was present in both cultures. Every second day (for 10 days), the parasitaemia was measured by flow cytometry, and the parasitised erythrocytes in both cultures were diluted by the same factor with uninfected erythrocytes (with the dilution yielding a 0.5% parasitaemia for the culture lacking GlcN).

### Western blots

Western blots were used to determine whether PfATP2-HA was successfully knocked down in PfATP2-HAreg parasites exposed to GlcN. Trophozoite-stage parasites were isolated from cultures that had been maintained with GlcN (5 mM for 7 days) or without GlcN. Parasite pellets (obtained by centrifuging the parasites at 12,000 x g for ∼ 30 s and removing the supernatant solution) were frozen at-80℃ then thawed. The following were then added to the (∼ 60 μL) sample: 40 μL 25 mM MgCl_2_, 1 μL 100x Protease Inhibitor Cocktail (Set III) (Merck Millipore), and 1 μL of Pierce^TM^ Universal Nuclease for Cell Lysis (250 units; ThermoFisher Scientific). The sample was vortexed then heated at 37°C for 5 min. NuPAGE® LDS Sample Buffer (1×; 100 µL) and 1× NuPAGE Sample Reducing Agent (40 µL) were then added, followed by 1× PBS to bring the sample volume up to 400 μL. The samples were run on NuPAGE Bis-Tris Mini Protein Gels (4-12%). After completion of electrophoresis, proteins in the gel were transferred to a nitrocellulose membrane following the NuPAGE Technical Guide (ThermoFisher Scientific). The membrane was then incubated in blocking buffer (4% w/v skim milk in 1× PBS) overnight at 4°C with constant shaking. The membranes were then exposed to specific primary and secondary antibodies (diluted in blocking buffer) for 1.5 h at room temperature in a humidified container. The primary antibodies used in this study were an anti-HA rat monoclonal antibody (1:500 dilution, clone 3F10, Roche) and an anti-heat shock protein 70 (HSP70) mouse monoclonal antibody (1:2000 dilution, a kind gift from Prof. Alex Maier). The secondary antibodies used were an anti-rat goat horseradish peroxidase (HRP)-conjugated antibody (1:5000 dilution, ab97057, Abcam) or an anti-mouse goat HRP-conjugated antibody (1:5000 dilution, ab6789, Abcam).

Subsequently, the membranes were washed with 1× PBS and visualised using Pierce™ ECL Plus Western Blotting Substrate with a ChemiDoc^TM^ MP Imaging System (Bio-Rad).

### Localisation of PfATP2-GFP in live cells

Erythrocytes infected with trophozoite-stage *P. falciparum* parasites were centrifuged (2,000 × *g*, 30 s) and resuspended in pH 7.4 Physiological Saline (125 mM NaCl, 5 mM KCl, 1 mM MgCl_2_, 20 mM glucose, 25 mM HEPES; pH 7.4) containing 20 μg/mL Hoechst 33258. After a 15 min incubation at 37℃, the cells were washed twice then resuspended in pH 7.4 Physiological Saline. An aliquot of the cell suspension was then added to the centre of a microscope slide, covered with a coverslip and sealed with nail polish. The cells were observed with an Applied Precision DeltaVision Elite system (GE Healthcare) with an inverted IX71 microscope with a 100X UPlanSApo oil immersion lens (Olympus). Images were taken using a Photometrics Cool SNAP HQ2 camera and deconvolved and adjusted for contrast and brightness using SoftWoRx Suite 2.0 software. GFP fluorescence was detected using wavelengths of 490 nm (excitation) and 525 nm (emission). Hoechst 33258 was detected at 350 nm (excitation) and 455 nm (emission). Images were processed using ImageJ software.

### Immunofluorescence assays

Immunofluorescence assays (IFAs) were performed with infected erythrocytes according to a published method (78). In brief, parasitised erythrocytes (on some occasions separated from uninfected erythrocytes using a Miltenyi Biotec VarioMACS magnet) were fixed with 4% paraformaldehyde and 0.0075% glutaraldehyde in 1×PBS for 30 min. After washing with 1×PBS, cells were permeabilised in 0.1% Triton X- 100 in 1×PBS for 10 min, treated with 0.1 M glycine in H_2_O for 15 min, then incubated overnight in 3% BSA in 1×PBS. Anti-HA High Affinity (1:200 dilution; clone 3F10, Roche) and rabbit anti-EXP2 antibody (1:500 dilution; (79)) were used for the primary antibody. After a 1 h incubation with primary antibody, cells were washed three times in 1×PBS and then incubated with Alexa Fluor 488 conjugated Donkey anti-Rat IgG (H+L) Highly Cross-Adsorbed Secondary Antibody (1:500 dilution; Catalog # A-21208, ThermoFisher Scientific) or Alexa Fluor 546 conjugated Goat anti-Rabbit IgG (H+L) Highly Cross-Adsorbed Secondary Antibody (1:400 dilution; Catalog # A-11035, ThermoFisher Scientific) for 1 h. The cells were treated with DAPI (Molecular Probes) for 5 min before being mounted on a slide. Microscopy was performed using an Applied Precision DeltaVision Elite system (as above) and images were processed using ImageJ.

### Measurements of parasite pH_cyt_

Parasite pH_cyt_ was measured under physiological conditions and in the presence of a low external [Cl^-^] using methods similar to those described previously (41,45). Briefly, isolated trophozoite-stage parasites suspended in bicarbonate-free medium were loaded with the pH-sensitive fluorescent dye BCECF by incubating them with BCECF-AM (5 μM) for 10 min at 37℃.

In the experiments for which data are shown in **Fig. 3**, the parasites were then washed twice in pH 7.1 Physiological Saline (125 mM NaCl, 5 mM KCl, 1 mM MgCl_2_, 20 mM glucose, 25 mM HEPES; pH 7.1) (with 12,000 × *g* 30 s centrifugation steps) and resuspended in this solution. The parasites were added to the compounds of interest dissolved in pH 7.1 Physiological Saline (or 0.1% v/v DMSO as the solvent control) in wells of a 96-well plate and fluorescence was recorded immediately.

In the experiments for which data are shown in **Fig. 4**, the parasites were washed and resuspended in Glucose-free Saline (135 mM NaCl, 5 mM KCl, 1 mM MgCl_2_, 25 mM HEPES; pH 7.1) after dye loading, and incubated at 37℃ for 20 min to allow for the depletion of ATP. The parasite suspension (15 μL per well) was then added to the compounds of interest dissolved in Glucose-free Cl^-^-free Saline (135 mM Na^+^- gluconate, 5 mM K^+^-gluconate, 1 mM MgSO_4_, 25 mM HEPES (pH 7.1)), yielding a final volume per well of 200 μL and an external [Cl^-^] of 11 mM, and fluorescence was recorded immediately.

For all pH_cyt_ experiments, fluorescence was measured at 37℃ with excitation wavelengths of 440 nm (yielding pH-insensitive fluorescence) and 495 nm (yielding pH-sensitive fluorescence). Emission was recorded at 520 nm.

To convert Fluorescence Ratios (495 nm/440 nm) into pH_cyt_ values, calibrations were included in all pH_cyt_ experiments. Parasites were suspended (at a density matching that used in the experimental wells) in calibration salines of known pH (130 mM KCl, 1 mM MgCl_2_, 20 mM glucose, 25 mM HEPES; pH 6.8, 7.1, 7.4, 7.8) containing nigericin (a compound that facilitates H^+^/K^+^ exchange; final concentration 5 μM), with fluorescence recorded for 5 min. The average of the Fluorescence Ratio values obtained during the read were plotted against the pH of the calibration solution, and a line was fit to the data. The resulting equation was used to convert Fluorescence Ratio values to pH_cyt_.

### Parasite proliferation assays

To evaluate the effects of compounds on the *in vitro* proliferation of parasites, a SYBR Safe-based fluorescence assay was used (80,81). The starting parasitaemia was 1% (with predominantly ring-stage parasites), the haematocrit was 1%, and the duration of the assay was 72 h. *P. falciparum*-infected erythrocytes incubated in the presence of 0.5 μM chloroquine were used as a non-proliferation control and those incubated in the absence of any compounds served as a 100% parasite proliferation control. The DMSO concentration in the assays did not exceed 0.1% (v/v). Parasite proliferation was measured and calculated as described previously (81). The parasite proliferation data were fitted with the equation Y = Bottom + (Top – Bottom)/(1 + (IC_50_/X)×Hillslope) using GraphPad Prism, where Y represents percentage parasite proliferation, IC_50_ represents the concentration of the test compound resulting in 50% inhibition of parasite proliferation, and X represents the compound concentration.

In the experiments for which data are shown in **Fig. S2**, the compounds were washed off 24 h into the 72 h experiment. Two washes were performed (with 1000 × g, 5 min centrifugation steps), bringing the concentration of compound in each well to ∼ 3% of its starting concentration.

### Phospholipid internalisation assays

For most experiments, phospholipid internalisation assays were performed with saponin-isolated trophozoite-stage parasites. After isolation, the parasites were washed three times in pH 7.1 Physiological Saline (37℃). In the case of parasites that had been exposed to 5 mM GlcN in the lead up to the experiments, 5 mM GlcN was present for the first two washes but not included in the third wash (and was absent for the remainder of the experiment). Parasites that were not exposed to GlcN were exposed to an equivalent volume of the solvent (complete medium lacking Albumax II; final concentration 1% v/v) during the relevant steps. For measurements performed with ATP-depleted parasites, parasites were washed and resuspended in Glucose-free Saline (containing 5 mM GlcN where appropriate), then incubated for ∼ 15 min at 37℃. The parasites were washed in Glucose-free Saline to remove GlcN prior to the NBD-PS uptake measurements. With the exception of the experiments for which data are shown in **Fig. S5** (in which parasites were maintained at 37℃ prior to, and during, the NBD-PS uptake measurements), all other measurements with isolated parasites were carried out at 15℃. Parasites (90 µL) were added to 100 µL of pH 7.1 Physiological Saline (or Glucose-free Saline for measurements with ATP-depleted parasites) (pre-warmed to 15℃) containing the compound of interest (or solvent alone). Unless stated otherwise, parasites were exposed to test compounds for 10 min while cooling to 15℃, before the addition of 10 µL NBD-PS (final concentration 5 μM). In all measurements, the final concentration of DMSO (used as a solvent for the MMV compounds) was 0.1% v/v, the concentration of water (solvent for vanadate) was 2% v/v, and the concentration of ethanol (solvent for NBD-PS) was 0.5% v/v. To terminate NBD-phospholipid uptake at the desired time point, an aliquot of cell suspension (150 μL) was transferred to a tube containing 1 mL ice-cold pH 7.1 Physiological Saline (or Glucose-free Saline for measurements with ATP-depleted parasites) containing 4% w/v fatty-acid free BSA. The samples were centrifuged (12,000 × g, 1 min, 4℃), the supernatant solution discarded, and the parasites resuspended in 1 mL of the same BSA-containing solution. The samples were centrifuged again, the supernatant solution discarded, and the parasites resuspended in 600 μL pH 7.1 Physiological Saline (or Glucose-free Saline in the case of ATP-depleted parasites) (37℃) containing the DNA stain Hoechst 33258 (2 μg/mL) for 15 min. The parasites were then washed and resuspended in 300 μL pH 7.1 Physiological Saline and maintained at room temperature until they were analysed by flow cytometry using a BD LSRII cytometer at the Cytometry, Histology and Spatial Multiomics Facility (Australian National University). The excitation wavelength was 488 nm for NBD-phospholipid fluorescence (530/30 nm emission filter), and 405 nm for Hoechst 33258 fluorescence (450/50 emission filter). Twenty thousand cells were sampled at low sampling speed with the following settings: forward scatter = 532 V (log scale), side scatter = 205 V (log scale), FITC = 852 V (log scale) and Pacific Blue = 418 V (log scale). The gating strategy is shown in **Fig. S10**. The geometric mean of NBD-phospholipid fluorescence for the population in the Hoechst 33258 positive, NBD-phospholipid positive gate was determined with FlowJo.

For experiments with erythrocytes (parasitised and uninfected) we adopted an approach described previously (49). Erythrocytes (5-10% of which were infected with trophozoite-stage parasites) were washed and resuspended in bicarbonate-free medium and exposed to the test compounds of interest for 10 min at 37℃. NBD-PS (final concentration 1 μM) and Hoechst 33258 (4 μg/mL) were then added and cells (final haematocrit 3.5-4.5%) were incubated at 37℃ for 30 min. The cells were then centrifuged (12,000 × g ∼ 30 s) and washed twice with 1 mL bicarbonate-free medium (37℃) containing 5% w/v fatty-acid free BSA. The cells were then resuspended in 1 mL 1×PBS and analysed by flow cytometry (as above).

### Microscopic determination of NBD-PS localisation

The location of NBD-PS was investigated in saponin-isolated trophozoite-stage parasites that had been exposed to 5 μM NBD-PS at 15°C in the dark for 9 min then washed with ice-cold 4% w/v BSA in pH 7.1 Physiological Saline (as above). Imaging was performed at the Centre for Advanced Microscopy (Australian National University) on a Leica STELLARIS 8, using a HC PL APO CS2 63x/1.40 OIL objective, with the Leica Lightning adaptive deconvolution module in the Leica Application Suite X (LAS-X) software (version 4.7.0.28176). The final fluorescent images were obtained using the wavelengths of 467 nm (excitation wavelength), 484 nm to 732 nm (emission wavelength), and transmitted light with Differential Interference Contrast (DIC).

## Acknowledgements

We are grateful to the Canberra Branch of the Australian Red Cross Lifeblood for the provision of blood and to the Medicines for Malaria Venture for the provision of MMV665794 and MMV007224. We also thank Michael Devoy (Cytometry, Histology and Spatial Multiomics Facility, The John Curtin School of Medical Research, Australian National University (ANU)) for assistance with flow cytometry, and Dr Teresa Neeman at the Biological Data Science Institute (ANU) for advice on statistical analyses. The authors acknowledge Microscopy Australia at the Centre for Advanced Microscopy, ANU, a facility enabled by NCRIS and university support.

## References

1. World Health Organization. (2023) World Malaria Report.

2. Rasmussen, C., Alonso, P., and Ringwald, P. (2022) Current and emerging strategies to combat antimalarial resistance. Expert Rev Anti Infect Ther 20, 353–372

3. Ward, K. E., Fidock, D. A., and Bridgford, J. L. (2022) *Plasmodium falciparum* resistance to artemisinin-based combination therapies. Curr Opin Microbiol 69, 102193

4. Cowell, A. N., Istvan, E. S., Lukens, A. K., Gomez-Lorenzo, M. G., Vanaerschot, M., Sakata-Kato, T., Flannery, E. L., Magistrado, P., Owen, E., Abraham, M., LaMonte, G., Painter, H. J., Williams, R. M., Franco, V., Linares, M., Arriaga, I., Bopp, S., Corey, V. C., Gnadig, N. F., Coburn-Flynn, O., Reimer, C., Gupta, P., Murithi, J. M., Moura, P. A., Fuchs, O., Sasaki, E., Kim, S. W., Teng, C. H., Wang, L. T., Akidil, A., Adjalley, S., Willis, P. A., Siegel, D., Tanaseichuk, O., Zhong, Y., Zhou, Y., Llinas, M., Ottilie, S., Gamo, F. J., Lee, M. C. S., Goldberg, D. E., Fidock, D. A., Wirth, D. F., and Winzeler, E. A. (2018) Mapping the malaria parasite druggable genome by using in vitro evolution and chemogenomics. Science 359, 191–199

5. Martin, R. E. (2020) The transportome of the malaria parasite. Biol Rev Camb Philos Soc 95, 305–332

6. Raj, D. K., Mu, J., Jiang, H., Kabat, J., Singh, S., Sullivan, M., Fay, M. P., McCutchan, T. F., and Su, X. Z. (2009) Disruption of a *Plasmodium falciparum* multidrug resistance-associated protein (PfMRP) alters its fitness and transport of antimalarial drugs and glutathione. J Biol Chem 284, 7687–7696

7. Rottmann, M., McNamara, C., Yeung, B. K., Lee, M. C., Zou, B., Russell, B., Seitz, P., Plouffe, D. M., Dharia, N. V., Tan, J., Cohen, S. B., Spencer, K. R., Gonzalez-Paez, G. E., Lakshminarayana, S. B., Goh, A., Suwanarusk, R., Jegla, T., Schmitt, E. K., Beck, H. P., Brun, R., Nosten, F., Renia, L., Dartois, V., Keller, T. H., Fidock, D. A., Winzeler, E. A., and Diagana, T. T. (2010) Spiroindolones, a potent compound class for the treatment of malaria. Science 329, 1175–1180

8. Spillman, N. J., Allen, R. J., McNamara, C. W., Yeung, B. K., Winzeler, E. A., Diagana, T. T., and Kirk, K. (2013) Na^+^ regulation in the malaria parasite *Plasmodium falciparum* involves the cation ATPase PfATP4 and is a target of the spiroindolone antimalarials. Cell Host Microbe 13, 227–237

9. Golldack, A., Henke, B., Bergmann, B., Wiechert, M., Erler, H., Blancke Soares, A., Spielmann, T., and Beitz, E. (2017) Substrate-analogous inhibitors exert antimalarial action by targeting the *Plasmodium* lactate transporter PfFNT at nanomolar scale. PLoS Pathog 13, e1006172

10. Hapuarachchi, S. V., Cobbold, S. A., Shafik, S. H., Dennis, A. S., McConville, M. J., Martin, R. E., Kirk, K., and Lehane, A. M. (2017) The malaria parasite’s lactate transporter PfFNT is the target of antiplasmodial compounds identified in whole cell phenotypic screens. PLoS Pathog 13, e1006180

11. Murithi, J. M., Deni, I., Pasaje, C. F. A., Okombo, J., Bridgford, J. L., Gnadig, N. F., Edwards, R. L., Yeo, T., Mok, S., Burkhard, A. Y., Coburn-Flynn, O., Istvan, E. S., Sakata-Kato, T., Gomez-Lorenzo, M. G., Cowell, A. N., Wicht, K. J., Le Manach, C., Kalantarov, G. F., Dey, S., Duffey, M., Laleu, B., Lukens, A. K., Ottilie, S., Vanaerschot, M., Trakht, I. N., Gamo, F. J., Wirth, D. F., Goldberg, D. E., Odom John, A. R., Chibale, K., Winzeler, E. A., Niles, J. C., and Fidock, D. A. (2022) The *Plasmodium falciparum* ABC transporter ABCI3 confers parasite strain-dependent pleiotropic antimalarial drug resistance. Cell Chem Biol 29, 824–839 e826

12. Martin, R. E., Shafik, S. H., and Richards, S. N. (2018) Mechanisms of resistance to the partner drugs of artemisinin in the malaria parasite. Curr Opin Pharmacol 42, 71–80

13. Richards, S. N., Nash, M. N., Baker, E. S., Webster, M. W., Lehane, A. M., Shafik, S. H., and Martin, R. E. (2016) Molecular mechanisms for drug hypersensitivity induced by the malaria parasite’s Chloroquine Resistance Transporter. PLoS Pathog 12, e1005725

14. Gupta, H. K., Shrivastava, S., and Sharma, R. (2017) A novel calcium uptake transporter of uncharacterized P-Type ATPase family supplies calcium for cell surface integrity in *Mycobacterium smegmatis*. mBio 8, e01388–17

15. Palmgren, M. (2023) P-type ATPases: Many more enigmas left to solve. J Biol Chem 299, 105352

16. Ashton, T., Dans, M., Favuzza, P., Ngo, A., Lehane, A. M., Zhang, X., Qiu, D., Chandra Maity, B., De, N., Schindler, K., Yeo, T., Park, H., Uhlemann, A., Churchyard, A., Baum, J., Fidock, D. A., Jarman, K., Lowes, K., Baud, D., Brand, S., Jackson, P., Cowman, A., and Sleebs, B. (2023) Optimization of 2,3-dihydroquinazolinone-3-carboxamides as antimalarials targeting PfATP4. J Med Chem 66, 3540–3565

17. Dennis, A. S. M., Rosling, J. E. O., Lehane, A. M., and Kirk, K. (2018) Diverse antimalarials from whole-cell phenotypic screens disrupt malaria parasite ion and volume homeostasis. Sci Rep 8, 8795

18. Flannery, E. L., McNamara, C. W., Kim, S. W., Kato, T. S., Li, F., Teng, C. H., Gagaring, K., Manary, M. J., Barboa, R., Meister, S., Kuhen, K., Vinetz, J. M., Chatterjee, A. K., and Winzeler, E. A. (2015) Mutations in the P-type cation-transporter ATPase 4, PfATP4, mediate resistance to both aminopyrazole and spiroindolone antimalarials. ACS Chem Biol 10, 413–420

19. Gilson, P. R., Kumarasingha, R., Thompson, J., Zhang, X., Penington, J. S., Kalhor, R., Bullen, H. E., Lehane, A. M., Dans, M. G., de Koning-Ward, T. F., Holien, J. K., Soares da Costa, T. P., Hulett, M. D., Buskes, M. J., Crabb, B. S., Kirk, K., Papenfuss, A. T., Cowman, A. F., and Abbott, B. M. (2019) A 4-cyano-3-methylisoquinoline inhibitor of *Plasmodium falciparum* growth targets the sodium efflux pump PfATP4. Sci Rep 9, 10292

20. Hewitt, S. N., Dranow, D. M., Horst, B. G., Abendroth, J. A., Forte, B., Hallyburton, I., Jansen, C., Baragana, B., Choi, R., Rivas, K. L., Hulverson, M. A., Dumais, M., Edwards, T. E., Lorimer, D. D., Fairlamb, A. H., Gray, D. W., Read, K. D., Lehane, A. M., Kirk, K., Myler, P. J., Wernimont, A., Walpole, C., Stacy, R., Barrett, L. K., Gilbert, I. H., and Van Voorhis, W. C. (2017) Biochemical and structural characterization of selective allosteric inhibitors of the *Plasmodium falciparum* drug target, prolyl-tRNA-synthetase. ACS Infect Dis 3, 34–44

21. Jimenez-Diaz, M. B., Ebert, D., Salinas, Y., Pradhan, A., Lehane, A. M., Myrand-Lapierre, M. E., O’Loughlin, K. G., Shackleford, D. M., Justino de Almeida, M., Carrillo, A. K., Clark, J. A., Dennis, A. S., Diep, J., Deng, X., Duffy, S., Endsley, A. N., Fedewa, G., Guiguemde, W. A., Gomez, M. G., Holbrook, G., Horst, J., Kim, C. C., Liu, J., Lee, M. C., Matheny, A., Martinez, M. S., Miller, G., Rodriguez-Alejandre, A., Sanz, L., Sigal, M., Spillman, N. J., Stein, P. D., Wang, Z., Zhu, F., Waterson, D., Knapp, S., Shelat, A., Avery, V. M., Fidock, D. A., Gamo, F. J., Charman, S. A., Mirsalis, J. C., Ma, H., Ferrer, S., Kirk, K., Angulo-Barturen, I., Kyle, D. E., DeRisi, J. L., Floyd, D. M., and Guy, R.K. (2014) (+)-SJ733, a clinical candidate for malaria that acts through ATP4 to induce rapid host-mediated clearance of *Plasmodium*. Proc Natl Acad Sci U S A 111, E5455–5462

22. Lehane, A. M., Ridgway, M. C., Baker, E., and Kirk, K. (2014) Diverse chemotypes disrupt ion homeostasis in the malaria parasite. Mol Microbiol 94, 327–339

23. Vaidya, A. B., Morrisey, J. M., Zhang, Z., Das, S., Daly, T. M., Otto, T. D., Spillman, N. J., Wyvratt, M., Siegl, P., Marfurt, J., Wirjanata, G., Sebayang, B. F., Price, R. N., Chatterjee, A., Nagle, A., Stasiak, M., Charman, S. A., Angulo-Barturen, I., Ferrer, S., Belen Jimenez-Diaz, M., Martinez, M. S., Gamo, F. J., Avery, V. M., Ruecker, A., Delves, M., Kirk, K., Berriman, M., Kortagere, S., Burrows, J., Fan, E., and Bergman, L. W. (2014) Pyrazoleamide compounds are potent antimalarials that target Na^+^ homeostasis in intraerythrocytic *Plasmodium falciparum*. Nat Commun 5, 5521

24. Best, J. T., Xu, P., and Graham, T. R. (2019) Phospholipid flippases in membrane remodeling and transport carrier biogenesis. Curr Opin Cell Biol 59, 8–15

25. Panatala, R., Hennrich, H., and Holthuis, J. C. (2015) Inner workings and biological impact of phospholipid flippases. J Cell Sci 128, 2021–2032

26. Norris, A. C., Mansueto, A. J., Jimenez, M., Yazlovitskaya, E. M., Jain, B. K., and Graham, T. R. (2024) Flipping the script: Advances in understanding how and why P4-ATPases flip lipid across membranes. Biochim Biophys Acta Mol Cell Res 1871, 119700

27. Lamy, A., Macarini-Bruzaferro, E., Dieudonne, T., Peralvarez-Marin, A., Lenoir, G., Montigny, C., le Maire, M., and Vazquez-Ibar, J. L. (2021) ATP2, The essential P4-ATPase of malaria parasites, catalyzes lipid-stimulated ATP hydrolysis in complex with a Cdc50 beta-subunit. Emerg Microbes Infect 10, 132–147

28. Perez-Victoria, F. J., Gamarro, F., Ouellette, M., and Castanys, S. (2003) Functional cloning of the miltefosine transporter. A novel P-type phospholipid translocase from *Leishmania* involved in drug resistance. J Biol Chem 278, 49965–49971

29. van Blitterswijk, W. J., and Verheij, M. (2013) Anticancer mechanisms and clinical application of alkylphospholipids. Biochim Biophys Acta 1831, 663–674

30. Wang, F., Li, X., Li, Y., Han, J., Chen, Y., Zeng, J., Su, M., Zhuo, J., Ren, H., Liu, H., Hou, L., Fan, Y., Yan, X., Song, S., Zhao, J., Jin, D., Zhang, M., and Pei, Y. (2021) *Arabidopsis* P4 ATPase-mediated cell detoxification confers resistance to *Fusarium graminearum* and *Verticillium dahliae*. Nat Commun 12, 6426

31. Zhang, M., Wang, C., Otto, T. D., Oberstaller, J., Liao, X., Adapa, S. R., Udenze, K., Bronner, I. F., Casandra, D., Mayho, M., Brown, J., Li, S., Swanson, J., Rayner, J. C., Jiang, R. H. Y., and Adams, J. H. (2018) Uncovering the essential genes of the human malaria parasite *Plasmodium falciparum* by saturation mutagenesis. Science 360, eaap7847

32. Cowan, N., Datwyler, P., Ernst, B., Wang, C., Vennerstrom, J. L., Spangenberg, T., and Keiser, J. (2015) Activities of N,N’-Diarylurea MMV665852 analogs against Schistosoma mansoni. Antimicrob Agents Chemother 59, 1935–1941

33. Debbert, S. L., Hintz, M. J., Bell, C. J., Earl, K. R., Forsythe, G. E., Haberli, C., and Keiser, J. (2021) Activities of Quinoxaline, Nitroquinoxaline, and [1,2,4]Triazolo[4,3-a]quinoxaline Analogs of MMV007204 against Schistosoma mansoni. Antimicrob Agents Chemother 65, e01370-20

34. Ingram-Sieber, K., Cowan, N., Panic, G., Vargas, M., Mansour, N. R., Bickle, Q. D., Wells, T. N., Spangenberg, T., and Keiser, J. (2014) Orally active antischistosomal early leads identified from the open access malaria box. PLoS Negl Trop Dis 8, e2610

35. Padalino, G., El-Sakkary, N., Liu, L. J., Liu, C., Harte, D. S. G., Barnes, R. E., Sayers, E., Forde-Thomas, J., Whiteland, H., Bassetto, M., Ferla, S., Johnson, G., Jones, A. T., Caffrey, C. R., Chalmers, I., Brancale, A., and Hoffmann, K. F. (2021) Anti-schistosomal activities of quinoxaline-containing compounds: From hit identification to lead optimisation. Eur J Med Chem 226, 113823

36. Prommana, P., Uthaipibull, C., Wongsombat, C., Kamchonwongpaisan, S., Yuthavong, Y., Knuepfer, E., Holder, A. A., and Shaw, P. J. (2013) Inducible knockdown of *Plasmodium* gene expression using the *glmS* ribozyme. PLoS One 8, e73783

37. de Souza, G. E., Bueno, R. V., de Souza, J. O., Zanini, C. L., Cruz, F. C., Oliva, G., Guido, R. V. C., and Aguiar, A. C. C. (2019) Antiplasmodial profile of selected compounds from Malaria Box: in vitro evaluation, speed of action and drug combination studies. Malar J 18, 447

38. Linares, M., Viera, S., Crespo, B., Franco, V., Gomez-Lorenzo, M. G., Jimenez-Diaz, M. B., Angulo-Barturen, I., Sanz, L. M., and Gamo, F. J. (2015) Identifying rapidly parasiticidal anti-malarial drugs using a simple and reliable in vitro parasite viability fast assay. Malar J 14, 441

39. Wu, J., Wang, C., Leas, D., Vargas, M., White, K. L., Shackleford, D. M., Chen, G., Sanford, A. G., Hemsley, R. M., Davis, P. H., Dong, Y., Charman, S. A., Keiser, J., and Vennerstrom, J. L. (2018) Progress in antischistosomal N,N’-diaryl urea SAR. Bioorg Med Chem Lett 28, 244–248

40. van Schalkwyk, D. A., Chan, X. W., Misiano, P., Gagliardi, S., Farina, C., and Saliba, K. J. (2010) Inhibition of *Plasmodium falciparum* pH regulation by small molecule indole derivatives results in rapid parasite death. Biochem Pharmacol 79, 1291–1299

41. Saliba, K. J., and Kirk, K. (1999) pH regulation in the intracellular malaria parasite, Plasmodium falciparum. H^+^ extrusion via a V-type H^+^-ATPase. J Biol Chem 274, 33213–33219

42. Saliba, K. J., and Kirk, K. (2001) H^+^-coupled pantothenate transport in the intracellular malaria parasite. J Biol Chem 276, 18115–18121

43. Alder, A., Sanchez, C. P., Russell, M. R. G., Collinson, L. M., Lanzer, M., Blackman, M. J., Gilberger, T. W., and Matz, J. M. (2023) The role of *Plasmodium* V-ATPase in vacuolar physiology and antimalarial drug uptake. Proc Natl Acad Sci U S A 120, e2306420120

44. Saliba, K. J., Allen, R. J., Zissis, S., Bray, P. G., Ward, S. A., and Kirk, K. (2003) Acidification of the malaria parasite’s digestive vacuole by a H^+^-ATPase and a H^+^-pyrophosphatase. J Biol Chem 278, 5605–5612

45. Henry, R. I., Cobbold, S. A., Allen, R. J., Khan, A., Hayward, R., Lehane, A. M., Bray, P. G., Howitt, S. M., Biagini, G. A., Saliba, K. J., and Kirk, K. (2010) An acid-loading chloride transport pathway in the intraerythrocytic malaria parasite, *Plasmodium falciparum*. J Biol Chem 285, 18615–18626

46. Lindblom, J. C. R., Zhang, X., and Lehane, A. M. (2024) A pH fingerprint assay to identify inhibitors of multiple validated and potential antimalarial drug targets. ACS Infect Dis 10, 1185–1200

47. Fraser, M., Jing, W., Broer, S., Kurth, F., Sander, L. E., Matuschewski, K., and Maier, A. G. (2021) Breakdown in membrane asymmetry regulation leads to monocyte recognition of *P. falciparum*-infected red blood cells. PLoS Pathog 17, e1009259

48. Calic, P. P. S., Ashton, T. D., Mansouri, M., Loi, K., Jarman, K. E., Qiu, D., Lehane, A. M., Roy, S., Rao, G. P., Maity, B., Wittlin, S., Crespo, B., Gamo, F. J., Deni, I., Fidock, D. A., Chowdury, M., de Koning-Ward, T. F., Cowman, A. F., Jackson, P. F., Baud, D., Brand, S., Laleu, B., and Sleebs, B. E. (2024) Optimization of pyrazolopyridine 4-carboxamides with potent antimalarial activity for which resistance is associated with the *P. falciparum* transporter ABCI3. Eur J Med Chem 276, 116677

49. Patel, A., Nofal, S. D., Blackman, M. J., and Baker, D. A. (2022) CDC50 orthologues in *Plasmodium falciparum* have distinct roles in merozoite egress and trophozoite maturation. mBio 13, e0163522

50. Chen, K., Gunay-Esiyok, O., Klingeberg, M., Marquardt, S., Pomorski, T. G., and Gupta, N. (2021) Aminoglycerophospholipid flipping and P4-ATPases in *Toxoplasma gondii*. J Biol Chem 296, 100315

51. Baldridge, R. D., and Graham, T. R. (2013) Two-gate mechanism for phospholipid selection and transport by type IV P-type ATPases. Proc Natl Acad Sci U S A 110, E358–367

52. Hiraizumi, M., Yamashita, K., Nishizawa, T., and Nureki, O. (2019) Cryo-EM structures capture the transport cycle of the P4-ATPase flippase. Science 365, 1149–1155

53. Timcenko, M., Dieudonne, T., Montigny, C., Boesen, T., Lyons, J. A., Lenoir, G., and Nissen, P. (2021) Structural basis of substrate-independent phosphorylation in a P4-ATPase lipid flippase. J Mol Biol 433, 167062

54. Xu, P., Baldridge, R. D., Chi, R. J., Burd, C. G., and Graham, T. R. (2013) Phosphatidylserine flipping enhances membrane curvature and negative charge required for vesicular transport. J Cell Biol 202, 875–886

55. Bisio, H., Krishnan, A., Marq, J. B., and Soldati-Favre, D. (2022) *Toxoplasma gondii* phosphatidylserine flippase complex ATP2B-CDC50.4 critically participates in microneme exocytosis. PLoS Pathog 18, e1010438

56. Chen, K., Huang, X., Distler, U., Tenzer, S., Gunay-Esiyok, O., and Gupta, N. (2023) Apically-located P4- ATPase1-Lem1 complex internalizes phosphatidylserine and regulates motility-dependent invasion and egress in *Toxoplasma gondii*. Comput Struct Biotechnol J 21, 1893–1906

57. Natarajan, P., Liu, K., Patil, D. V., Sciorra, V. A., Jackson, C. L., and Graham, T. R. (2009) Regulation of a Golgi flippase by phosphoinositides and an ArfGEF. Nat Cell Biol 11, 1421–1426

58. Kilian, N., Choi, J. Y., Voelker, D. R., and Ben Mamoun, C. (2018) Role of phospholipid synthesis in the development and differentiation of malaria parasites in the blood. J Biol Chem 293, 17308–17316

59. Wein, S., Ghezal, S., Bure, C., Maynadier, M., Perigaud, C., Vial, H. J., Lefebvre-Tournier, I., Wengelnik, K., and Cerdan, R. (2018) Contribution of the precursors and interplay of the pathways in the phospholipid metabolism of the malaria parasite. J Lipid Res 59, 1461–1471

60. Nofal, S. D., Patel, A., Blackman, M. J., Flueck, C., and Baker, D. A. (2021) *Plasmodium falciparum* guanylyl cyclase-Alpha and the activity of its appended P4-ATPase domain are essential for cGMP synthesis and blood-stage egress. mBio 12, e02694–20

61. Yang, Z., Shi, Y., Cui, H., Yang, S., Gao, H., and Yuan, J. (2021) A malaria parasite phospholipid flippase safeguards midgut traversal of ookinetes for mosquito transmission. Sci Adv 7, eabf6015

62. Taylor, C. J., McRobert, L., and Baker, D. A. (2008) Disruption of a *Plasmodium falciparum* cyclic nucleotide phosphodiesterase gene causes aberrant gametogenesis. Mol Microbiol 69, 110–118

63. Haase, S., Condron, M., Miller, D., Cherkaoui, D., Jordan, S., Gulbis, J. M., and Baum, J. (2021) Identification and characterisation of a phospholipid scramblase in the malaria parasite *Plasmodium falciparum*. Mol Biochem Parasitol 243, 111374

64. Cheng, T., Zhao, Y., Li, X., Lin, F., Xu, Y., Zhang, X., Li, Y., Wang, R., and Lai, L. (2007) Computation of octanol-water partition coefficients by guiding an additive model with knowledge. J Chem Inf Model 47, 2140–2148

65. Kumpornsin, K., Kochakarn, T., Yeo, T., Okombo, J., Luth, M. R., Hoshizaki, J., Rawat, M., Pearson, R. D., Schindler, K. A., Mok, S., Park, H., Uhlemann, A. C., Jana, G. P., Maity, B. C., Laleu, B., Chenu, E., Duffy, J., Moliner Cubel, S., Franco, V., Gomez-Lorenzo, M. G., Gamo, F. J., Winzeler, E. A., Fidock, D. A., Chookajorn, T., and Lee, M. C. S. (2023) Generation of a mutator parasite to drive resistome discovery in *Plasmodium falciparum*. Nat Commun 14, 3059

66. Van Voorhis, W. C., Adams, J. H., Adelfio, R., Ahyong, V., Akabas, M. H., Alano, P., Alday, A., Aleman Resto, Y., Alsibaee, A., Alzualde, A., Andrews, K. T., Avery, S. V., Avery, V. M., Ayong, L., Baker, M., Baker, S., Ben Mamoun, C., Bhatia, S., Bickle, Q., Bounaadja, L., Bowling, T., Bosch, J., Boucher, L. E., Boyom, F. F., Brea, J., Brennan, M., Burton, A., Caffrey, C. R., Camarda, G., Carrasquilla, M., Carter, D., Belen Cassera, M., Chih-Chien Cheng, K., Chindaudomsate, W., Chubb, A., Colon, B. L., Colon-Lopez, D. D., Corbett, Y., Crowther, G. J., Cowan, N., D’Alessandro, S., Le Dang, N., Delves, M., DeRisi, J. L., Du, A. Y., Duffy, S., Abd El-Salam El-Sayed, S., Ferdig, M. T., Fernandez Robledo, J. A., Fidock, D. A., Florent, I., Fokou, P. V., Galstian, A., Gamo, F. J., Gokool, S., Gold, B., Golub, T., Goldgof, G. M., Guha, R., Guiguemde, W. A., Gural, N., Guy, R. K., Hansen, M. A., Hanson, K. K., Hemphill, A., Hooft van Huijsduijnen, R., Horii, T., Horrocks, P., Hughes, T. B., Huston, C., Igarashi, I., Ingram-Sieber, K., Itoe, M. A., Jadhav, A., Naranuntarat Jensen, A., Jensen, L. T., Jiang, R. H., Kaiser, A., Keiser, J., Ketas, T., Kicka, S., Kim, S., Kirk, K., Kumar, V. P., Kyle, D. E., Lafuente, M. J., Landfear, S., Lee, N., Lee, S., Lehane, A. M., Li, F., Little, D., Liu, L., Llinas, M., Loza, M. I., Lubar, A., Lucantoni, L., Lucet, I., Maes, L., Mancama, D., Mansour, N. R., March, S., McGowan, S., Medina Vera, I., Meister, S., Mercer, L., Mestres, J., Mfopa, A. N., Misra, R. N., Moon, S., Moore, J. P., Morais Rodrigues da Costa, F., Muller, J., Muriana, A., Nakazawa Hewitt, S., Nare, B., Nathan, C., Narraidoo, N., Nawaratna, S., Ojo, K. K., Ortiz, D., Panic, G., Papadatos, G., Parapini, S., Patra, K., Pham, N., Prats, S., Plouffe, D. M., Poulsen, S. A., Pradhan, A., Quevedo, C., Quinn, R. J., Rice, C. A., Abdo Rizk, M., Ruecker, A., St Onge, R., Salgado Ferreira, R., Samra, J., Robinett, N. G., Schlecht, U., Schmitt, M., Silva Villela, F., Silvestrini, F., Sinden, R., Smith, D. A., Soldati, T., Spitzmuller, A., Stamm, S. M., Sullivan, D. J., Sullivan, W., Suresh, S., Suzuki, B. M., Suzuki, Y., Swamidass, S. J., Taramelli, D., Tchokouaha, L. R., Theron, A., Thomas, D., Tonissen, K. F., Townson, S., Tripathi, A. K., Trofimov, V., Udenze, K. O., Ullah, I., Vallieres, C., Vigil, E., Vinetz, J. M., Voong Vinh, P., Vu, H., Watanabe, N. A., Weatherby, K., White, P. M., Wilks, A. F., Winzeler, E. A., Wojcik, E., Wree, M., Wu, W., Yokoyama, N., Zollo, P. H., Abla, N., Blasco, B., Burrows, J., Laleu, B., Leroy, D., Spangenberg, T., Wells, T., and Willis, P. A. (2016) Open source drug discovery with the Malaria Box compound collection for neglected diseases and beyond. PLoS Pathog 12, e1005763

67. Arashiki, N., Takakuwa, Y., Mohandas, N., Hale, J., Yoshida, K., Ogura, H., Utsugisawa, T., Ohga, S., Miyano, S., Ogawa, S., Kojima, S., and Kanno, H. (2016) ATP11C is a major flippase in human erythrocytes and its defect causes congenital hemolytic anemia. Haematologica 101, 559–565

68. Bryk, A. H., and Wisniewski, J. R. (2017) Quantitative analysis of human red blood cell proteome. J Proteome Res 16, 2752–2761

69. Liou, A. Y., Molday, L. L., Wang, J., Andersen, J. P., and Molday, R. S. (2019) Identification and functional analyses of disease-associated P4-ATPase phospholipid flippase variants in red blood cells. J Biol Chem 294, 6809–6821

70. Rawat, M., Padalino, G., Yeo, T., Brancale, A., Fidock, D. A., Hoffmann, K. F., and Lee, M. C. S. (2024) Quinoxaline-based anti-schistosomal compounds have potent anti-malarial activity. bioRxiv

71. Rug, M., and Maier, A. G. (2013) Transfection of *Plasmodium falciparum*. Methods Mol Biol 923, 75–98

72. McHugh, E., Batinovic, S., Hanssen, E., McMillan, P. J., Kenny, S., Griffin, M. D., Crawford, S., Trenholme, K. R., Gardiner, D. L., Dixon, M. W., and Tilley, L. (2015) A repeat sequence domain of the ring-exported protein-1 of *Plasmodium falciparum* controls export machinery architecture and virulence protein trafficking. Mol Microbiol 98, 1101–1114

73. Lambros, C., and Vanderberg, J. P. (1979) Synchronization of *Plasmodium falciparum* erythrocytic stages in culture. J Parasitol 65, 418–420

74. Wu, Y., Sifri, C. D., Lei, H. H., Su, X. Z., and Wellems, T. E. (1995) Transfection of *Plasmodium falciparum* within human red blood cells. Proc Natl Acad Sci U S A 92, 973–977

75. Qiu, D., Pei, J. V., Rosling, J. E. O., Thathy, V., Li, D., Xue, Y., Tanner, J. D., Penington, J. S., Aw, Y. T. V., Aw, J. Y. H., Xu, G., Tripathi, A. K., Gnadig, N. F., Yeo, T., Fairhurst, K. J., Stokes, B. H., Murithi, J. M., Kumpornsin, K., Hasemer, H., Dennis, A. S. M., Ridgway, M. C., Schmitt, E. K., Straimer, J., Papenfuss, A. T., Lee, M. C. S., Corry, B., Sinnis, P., Fidock, D. A., van Dooren, G. G., Kirk, K., and Lehane, A. M. (2022) A G358S mutation in the *Plasmodium falciparum* Na^+^ pump PfATP4 confers clinically-relevant resistance to cipargamin. Nat Commun 13, 5746

76. Livak, K. J., and Schmittgen, T. D. (2001) Analysis of relative gene expression data using real-time quantitative PCR and the 2(-Delta Delta C(T)) Method. Methods 25, 402–408

77. Burda, P. C., Crosskey, T., Lauk, K., Zurborg, A., Sohnchen, C., Liffner, B., Wilcke, L., Pietsch, E., Strauss, J., Jeffries, C. M., Svergun, D. I., Wilson, D. W., Wilmanns, M., and Gilberger, T. W. (2020) Structure-based identification and functional characterization of a lipocalin in the malaria parasite *Plasmodium falciparum*. Cell Rep 31, 107817

78. Tonkin, C. J., van Dooren, G. G., Spurck, T. P., Struck, N. S., Good, R. T., Handman, E., Cowman, A. F., and McFadden, G. I. (2004) Localization of organellar proteins in *Plasmodium falciparum* using a novel set of transfection vectors and a new immunofluorescence fixation method. Mol Biochem Parasitol 137, 13–21

79. Lawrence, N., Dennis, A. S. M., Lehane, A. M., Ehmann, A., Harvey, P. J., Benfield, A. H., Cheneval, O., Henriques, S. T., Craik, D. J., and McMorran, B. J. (2018) Defense peptides engineered from Human Platelet Factor 4 kill *Plasmodium* by selective membrane disruption. Cell Chem Biol 25, 1140–1150 e1145

80. Smilkstein, M., Sriwilaijaroen, N., Kelly, J. X., Wilairat, P., and Riscoe, M. (2004) Simple and inexpensive fluorescence-based technique for high-throughput antimalarial drug screening. Antimicrob Agents Chemother 48, 1803–1806

81. Spry, C., Macuamule, C., Lin, Z., Virga, K. G., Lee, R. E., Strauss, E., and Saliba, K. J. (2013) Pantothenamides are potent, on-target inhibitors of *Plasmodium falciparum* growth when serum pantetheinase is inactivated. PLoS One 8, e54974

