## Supporting information for "PfATP2 is an essential flippase on the *Plasmodium falciparum* surface that influences parasite sensitivity to antiplasmodial compounds"

***Table S1.*** *Primers used in this study. Restriction enzyme sites are underlined.*

| Primer name in study | Primer name in lab | Sequence (5' – 3') | Use |
|---|---|---|---|
| P1 | PfATP2 F1 | TTACATATAACTCGAGATGTCCTTAGTTTATAGAAAAACACTG AAC | Generation of 3D7-PfATP2+ |
| P2 | PfATP2 R2 | TCTTCTCCTTTACTGGTACCTTATTAAATCATATTATCTTGTTTT CTAATTAATTGTATA | Generation of 3D7-PfATP2+ |
| P3 | PfATP2 F4 | CGAGATCTACCTGTTGTAATTCATGCTGTGT | Generation of PfATP2-HAreg and PfATP2-GFPreg, sequencing for 3D7-PfATP2+ and PfATP2-HAreg vector |
| P4 | PfATP2 R4c | CGCTGCAGAATCATATTATCTTGTTTTCTAATTAATTGTATACA TG | Generation of PfATP2-HAreg |
| P5 | PfATP2 R4 | CGGGTACCAATCATATTATCTTGTTTTCTAATTAATTGTATACA TG | Generation of PfATP2-GFPreg |
| P6 | PfATP2 F7 | ATAATGAAGGGCGCACACCA | qRT-PCR and sequencing of *pfatp2* overexpression vector |
| P7 | PfATP2 R7 | TCTACATCGGTTCGTTTGGCT | qRT-PCR and sequencing of *pfatp2* overexpression vector |
| P8 | PfATP2 F8 | GGTGATGGAGCAAATGACCG | qRT-PCR, sequencing of *pfatp2* overexpression vector and integration checks (PfATP2-HAreg and PfATP2-GFPreg) |
| P9 | PfATP2 R8 | GCGGAATTGACTAATACCATAATCTGA | qRT-PCR and sequencing of *pfatp2* overexpression vector |
| P10 | 18S rRNA control F1 | GCTGACTACGTCCCTGCCC | qRT-PCR |
| P11 | 18S rRNA control R1 | ACAATTCATCATATCTTTCAATCGGTA | qRT-PCR |
| P12 | 18S rRNA control F2 | GGCAACAACAGGTCTGTGAT | qRT-PCR |
| P13 | 18S rRNA control R2 | TTCGGCGGAGGAAAAGTATG | qRT-PCR |
| P14 | stRNA F | AAGTAGCAGGTCATCGTGGTT | qRT-PCR |
| P15 | stRNA R | AGTTCGGCACATTCTTCCATAA | qRT-PCR |
| P16 | PfATP2 F5 | TTCAGAGGTCTCTATAG AATGTGTTATGGTATTGTAAGTGAAGAAG | sequencing for 3D7-PfATP2+ |
| P17 | PfATP2 R5 | AGCGTGGGTCTCGTACT ATGGTTCCTAAAAGATGAATACTTCCTAC | sequencing for 3D7-PfATP2+ |
| P18 | PfATP2 F6 | TGGGTAGTTATTACAGGAAATTTCGT | sequencing for 3D7-PfATP2+ |
| P19 | PfATP2 R6 | TCACCAAATGAAGGAACATCTGC | sequencing for 3D7-PfATP2+ |
| P20 | PfATP2f6 4 | AAAATCGATTTAGATATTTGCCTT | sequencing for 3D7-PfATP2+ |

| | | | |
|---|---|---|---|
| P21 | PfATP2r65 | TTCATGGTTGGTTCCACAGG | sequencing for 3D7-PfATP2+ |
| P22 | pGLUX Fwd | TCCGTTAATAATAAATACACGCAGTC | sequencing for 3D7-PfATP2+ |
| P23 | pGLUX rvs | TGTGCCCATTAACATCACCATC | sequencing for 3D7-PfATP2+ |
| P24 | M13R | CAGGAAACAGCTATGAC | sequencing for PfATP2-HAreg vector |
| P25 | Sp6 | TATTTAGGTGACACTATAG | sequencing for PfATP2-HAreg vector |
| P26 | glmsHA rvs | GTTTGAAGAAATCCTTACGGCTGTG | sequencing for PfATP2-HAreg vector; transfection and integration checks (PfATP2-HAreg) |
| P27 | glmsGFP rvs | GTAAGTTTTCCGTATGTTGCATCACC | sequencing for pGFP_*glmS* vector; transfection and integration checks (PfATP2-GFPreg) |
| P28 | GFPglms fwd | ATTATATTTTTTTCTTCCCACATTTCGT | sequencing for pGFP_*glmS* vector |
| P29 | PfATP2 r69 | TCAATCCATAGGATATATGATATGTAAGT | Paired with P8 to detect unmodified *pfatp2* locus |
| P30 | PfATP2 F3 | GATCGCGGCCGCTAACCATACGACATTTGGACCGT | Checks for successful transfection (PfATP2-HAreg and PfATP2-GFPreg) |

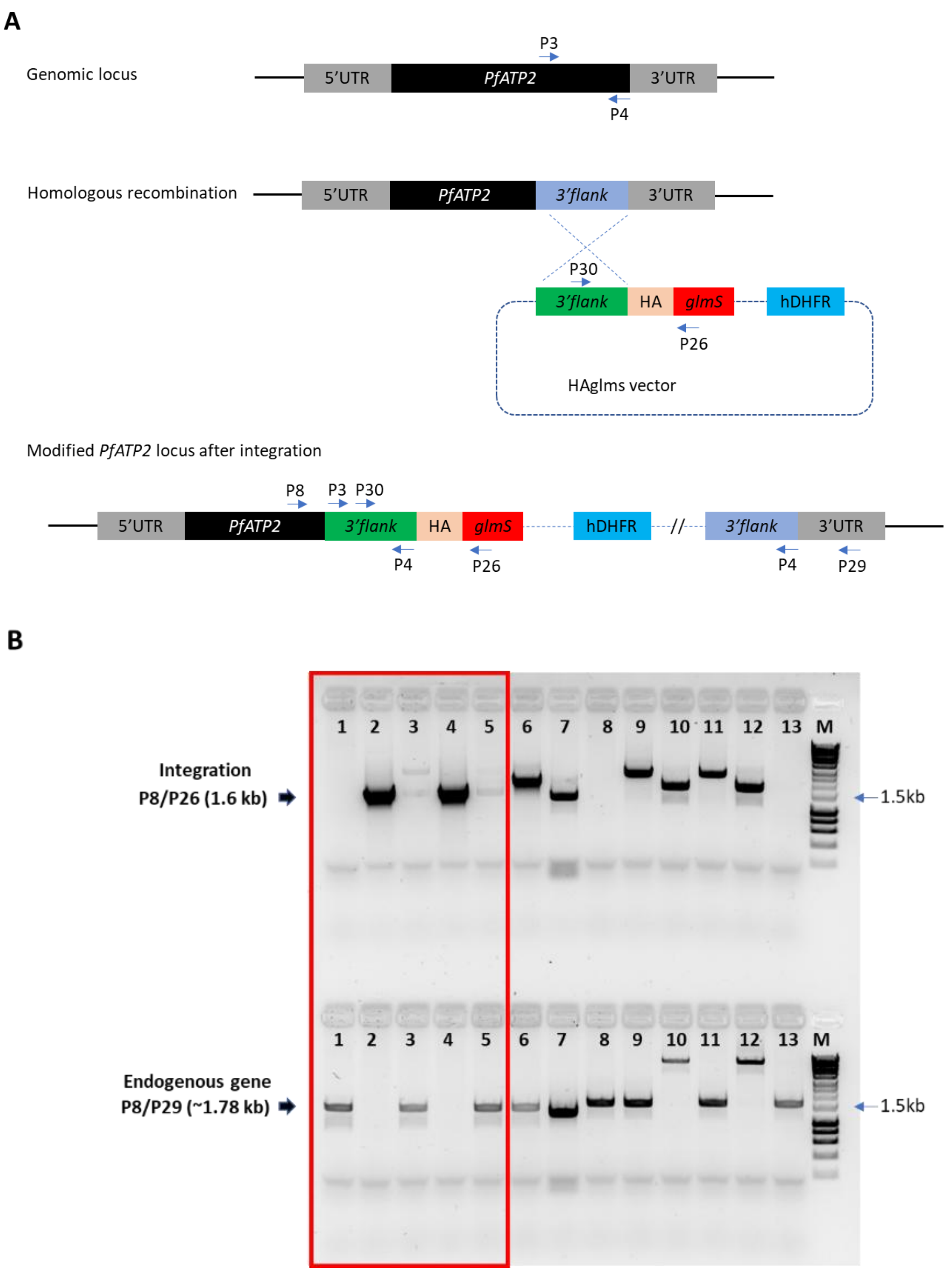


***Figure S1. Integration of the PfATP2-HA-glmS targeting construct into the endogenous* pfatp2 *locus.*** *(**A**) Schematic showing the strategy used to incorporate the HA and glmS ribozyme sequences at the 3' end of the* pfatp2 *gene by single crossover homologous recombination. The numbered arrows show the primers that were used to make the construct and to test for successful integration. P3 and P4 were used to amplify the 3' flank of* pfatp2*. Diagnostic PCRs to test for successful transfection and integration of the PfATP2-HA-glmS construct used primer pairs P30/P26, P8/P26, and P8/P29. UTR, untranslated region; HA,* 3x *Haemagglutinin epitope tag; glmS, glmS ribozyme sequence; hDHFR, human dihydrofolate reductase (selectable marker). (**B**) Image of PCR products run on an agarose gel. Primers P8 and P26 were used to test for integration of the desired DNA (expected size 1.6 kb) and primers P8 and P29 were used to test for the presence of the unmodified* pfatp2 *gene (expected size ~1.78 kb). Lane 1: PCRs performed with wild-type 3D7 parasites. Lanes 2 and 4: PCRs performed with PfATP2-HAreg clone F11 (the clone used throughout the study). Lanes 3 and 5: PCRs performed with PfABCI3-HAreg parasites. The contents of the other lanes are not related to this study. The lane labelled M was loaded with Hyperladder$^{TM}$ 1 kb (Bioline).*

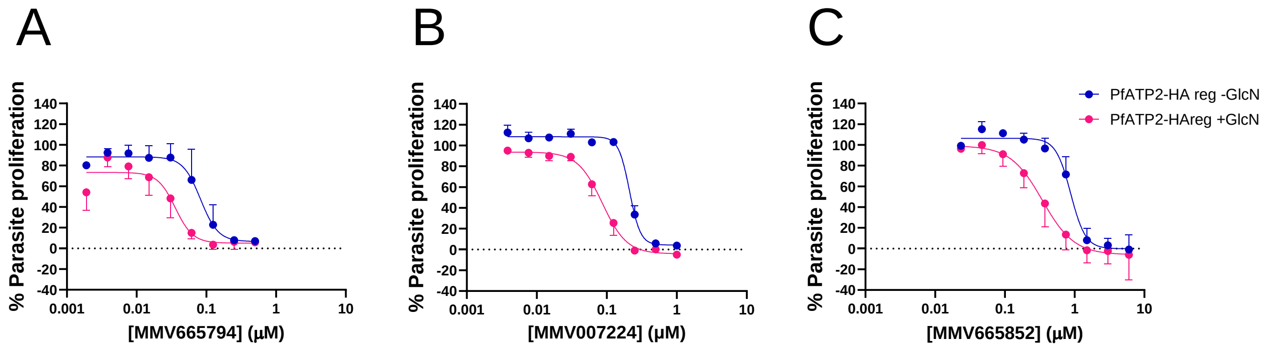


***Figure S2. Growth inhibition of PfATP2-HA knockdown and Control parasites after a 24 h exposure to MMV665794 (A), MMV007224 (B) and MMV665852 (C).*** *The data for parasites in which PfATP2-HA was knocked down (PfATP2-HAreg +GlcN) are shown in pink and the data for PfATP2-HAreg parasites that were not exposed to GlcN (-GlcN) are shown in blue. Where present, GlcN (5 mM) was added to cultures four days before the start of the experiment and maintained throughout the experiment. The duration of the assays was 72 h, with the compounds washed off 24 h after the start of the assay (see Materials and Methods). The data are from three independent experiments performed on different days, except for the highest and lowest concentrations, for which data are n = 1-2. The* $IC_{50}$ *values were 82 ± 25 nM (-GlcN) and 34.5 ± 10.0 nM (+GlcN) (mean ± SEM, n = 3,* P *= 0.03, ratio paired t-test) for MMV665794; 220 ± 12 nM (-GlcN) and 90 ± 21 nM (+GlcN) (mean ± SEM, n = 3,* P *= 0.05, ratio paired t-test) for MMV007224; and 852 ± 183 nM (-GlcN) and 343 ± 70 nM (+GlcN) (mean ± SEM, n = 3,* P *= 0.002, ratio paired t-test) for MMV665852.*

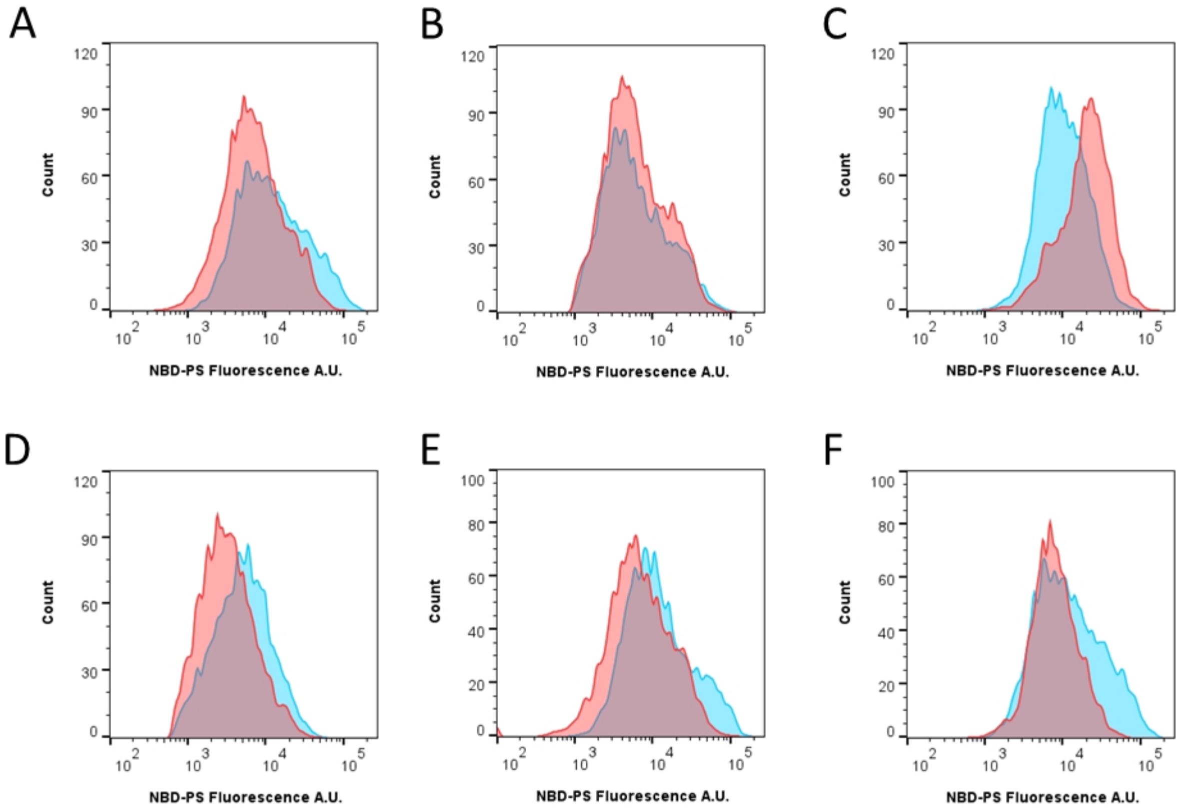


**Figure S3. Flow cytometry analysis of NBD-PS uptake in isolated trophozoites.** *NBD-PS internalisation was measured in isolated trophozoite-stage parasites suspended in pH 7.1 Physiological Saline at 15˚C. In all panels, the parasites were incubated with either a compound (**D**- **F**) or solvent alone (0.1% v/v DMSO; **A**-**C**) for 10 min, then NBD-PS uptake was measured over 9 min. Cells were gated for Hoechst 33258 fluorescence and NBD-PS fluorescence using the strategy shown in **Fig. S10**. In each panel, data from a single experiment, representative of at least three independent experiments, are shown. NBD-PS internalisation was measured in: **(A)** PfATP2-HAreg parasites in which PfATP2-HA was knocked down (red; two day exposure of culture to 5 mM GlcN) or expressed at a normal level (-GlcN; blue); (**B**) wild-type 3D7 parasites from cultures that were either exposed to 5 mM GlcN for two days in the lead-up to the experiment (red) or that were not exposed to GlcN (blue); (**C**)* pfatp2*-overexpressing 3D7-PfATP2+ parasites (red) and empty vector control parasites (3D7-EV; blue); **(D-F)** PfATP2-HAreg Control parasites (-GlcN) in the absence (blue) or presence (red) of (**D**) 10 µM MMV665794, (**E**) 15 µM MMV007224 (red) or **(F)** 500 µM vanadate.*

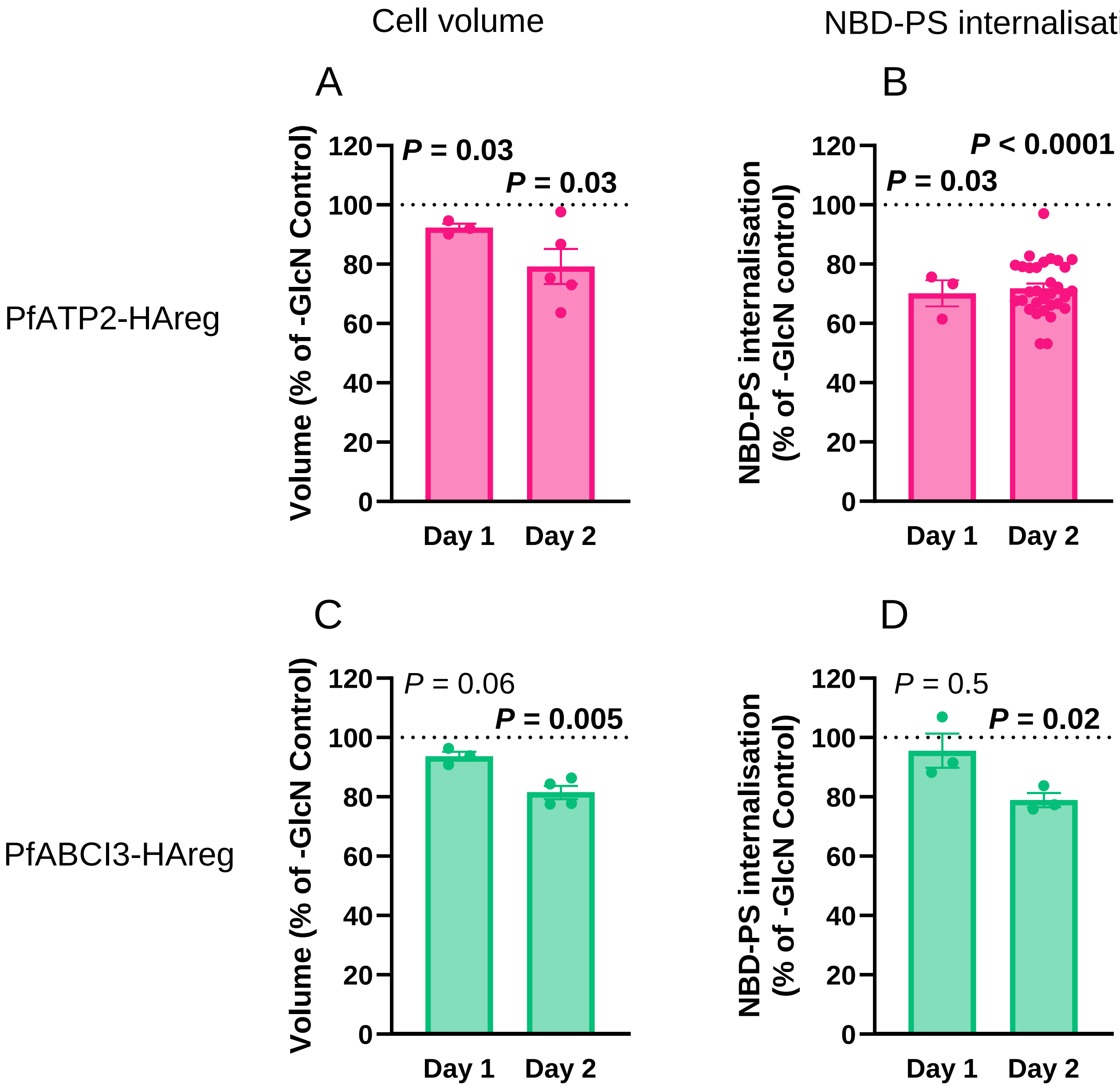


***Fig. S4. The effects of exposing PfATP2-HAreg or PfABCI3-HAreg parasites to GlcN for one or two days on parasite volume and NBD-PS internalisation.*** *(**A**,**C**) The mean volume of isolated trophozoite-stage PfATP2-HAreg (**A**) or PfABCI3-HAreg (**C**) parasites from cultures that had been exposed to 5 mM GlcN for one day or two days in the lead up to the experiment, expressed as a percentage of those obtained for the equivalent parasites that had not been exposed to GlcN. The* P *values are the results of ratio paired t-tests (with comparisons made to the -GlcN Control) performed with the pre-normalised data (volume in fL). The volumes measured in the different experiments ranged from 19-59 fL. (**B**,**D**) NBD-PS internalisation (measured over 9 min at 15˚C) by isolated trophozoite-stage PfATP2-HAreg (**B**) or PfABCI3-HAreg (**D**) parasites that had been exposed to 5 mM GlcN for one day or two days, expressed as a percentage of that observed for the equivalent parasites that had not been exposed to GlcN. The* P *values show the results of ratio paired t-tests performed on pre-normalised data (NBD-PS fluorescence (geometric mean), which ranged from 2183-24602 in different experiments). The Day 2 data in panel **B** summarise data from all the different experiments (n=31) performed in this study in which NBD-PS internalisation was measured for 9 min at 15˚C in isolated ATP-replete PfATP2-HAreg parasites from cultures that were either not exposed to GlcN or exposed to 5 mM GlcN for two days, drawing together data from **Fig. 5A**, **5D**, **6A**, **6B**, **S5B**, **S7**, **S8A** and **S8B**, **S9B** and **S9C**. In all panels, the bars and error bars show the mean and SEM, and the symbols show the data from individual biological replicates (performed on different days).* P *values indicating statistical significance (≤ 0.05) are shown in bold. GlcN was not present during the measurements.*

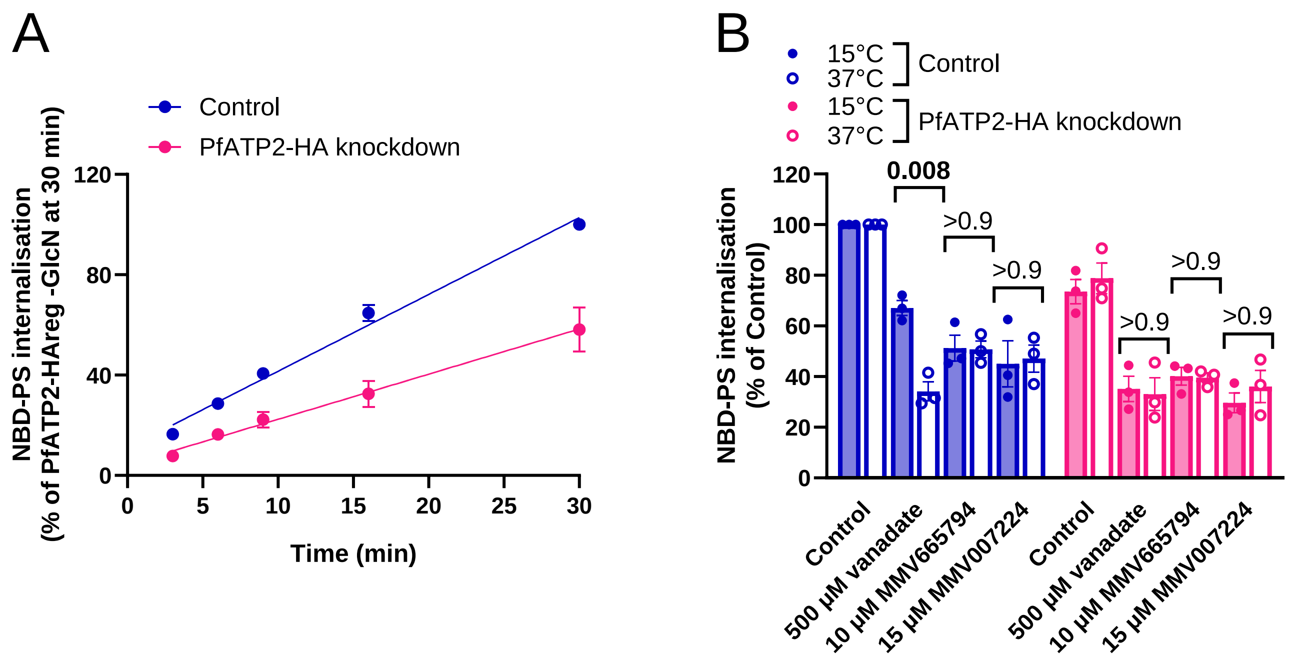


***Fig. S5. The effects of PfATP2-HA knockdown, vanadate, MMV665794 and MMV007224 on NBD-PS uptake by isolated trophozoite-stage parasites at 37˚C.*** *Both panels show data for PfATP2-HAreg parasites that were either not exposed to GlcN (Control; blue) or that were exposed to 5 mM GlcN for two days in the lead up to the experiment to reduce PfATP2-HA expression (PfATP2-HA knockdown; pink). GlcN was not present during the measurements. (**A**) The parasites were suspended in pH 7.1 Physiological Saline at 37˚C, with NBD-PS uptake measured at the time points indicated. (**B**) The parasites were suspended at pH 7.1 Physiological Saline at either 37˚C or 15˚C, in the absence (solvent control) or presence of the compounds and concentrations indicated, with NBD-PS internalisation measured at the 9 min time point. In **A**, the data shown are the mean ± SEM; in **B**, the symbols show the data from individual experiments, and the bars and error bars show the mean ± SEM. In both panels, the data are from three independent experiments (performed on different days; with all conditions tested concurrently) and are expressed as a percentage of the NBD-PS fluorescence (geometric mean) measured under the conditions indicated on the y axis. In **B**, the NBD-PS fluorescence (geometric mean) was higher at 37˚C (ranging from 19784-55878 for the solvent control -GlcN parasites in the different experiments) than at 15˚C (range: 5021-11144). The numbers above the bars are* P *values from a three-way ANOVA with post hoc Tukey test performed on the normalised data.*

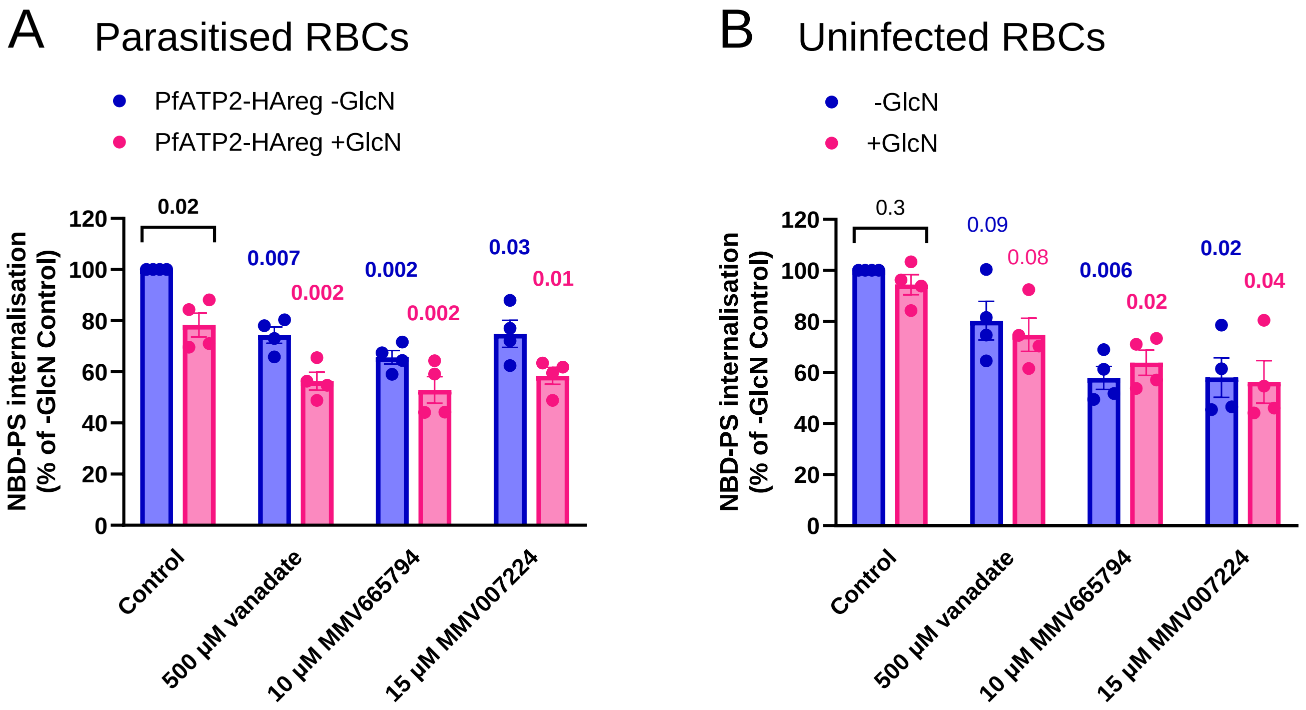


***Fig. S6. Effect of GlcN, vanadate (500 µM), MMV665794 (10 µM) and MMV007224 (15 µM) on NBD-PS internalisation by erythrocytes infected with PfATP2-HAreg parasites (A) and uninfected erythrocytes (B).*** *NBD-PS internalisation was measured over 30 min in cells suspended in bicarbonate-free medium at 37˚C. The cells were either exposed to 5 mM GlcN for two days in the lead up to the experiment to reduce PfATP2-HA expression (+GlcN; pink) or not exposed to GlcN (-GlcN; blue). The bars and error bars show the mean ± SEM (from four independent experiments performed on different days, with all conditions tested concurrently) and the symbols show the data from individual experiments. The data are expressed as a percentage of the NBD-PS fluorescence (geometric mean) measured in the -GlcN Control. The NBD-PS fluorescence for the -GlcN Control (geometric mean) was higher in parasitised erythrocytes (ranging from 2406 in experiment 1 (lowest) to 52323 in experiment 3 (highest)) than in uninfected erythrocytes (ranging from 795 in experiment 1 to 9143 in experiment 3). Statistical comparisons were made on the natural logarithm transformed pre-normalised data (NBD-PS fluorescence (geometric mean)) using paired t-tests. The* P *values are shown, with values indicating statistical significance (*P *≤ 0.05) shown in bold. Coloured values are the* P *values for comparisons with the solvent control for the same cell type.*

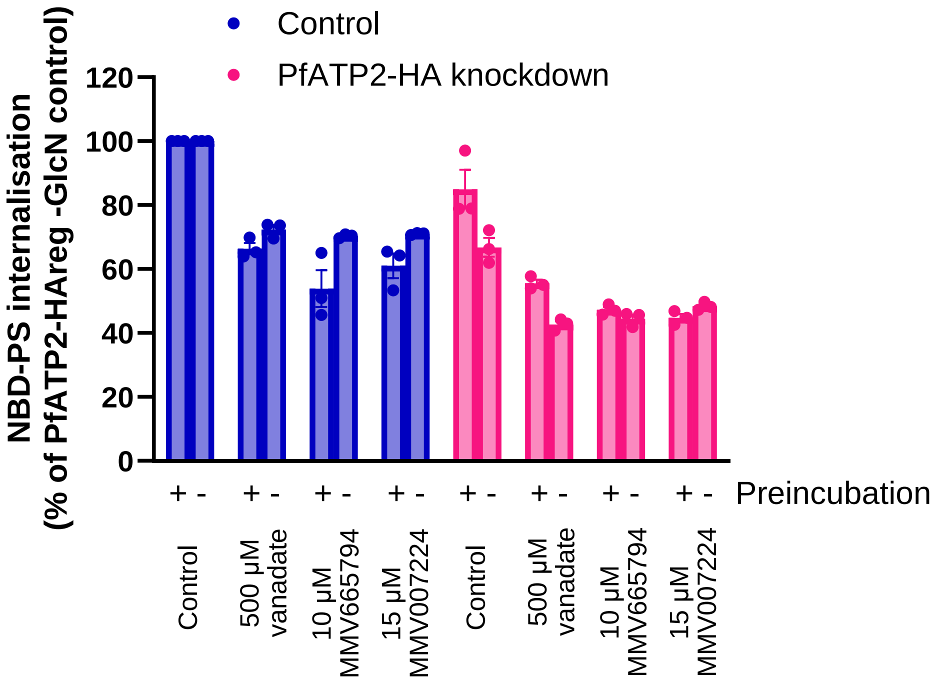


***Fig. S7. Effect of vanadate (500 μM), MMV665794 (10 μM) and MMV007224 (15 μM) on NBD-PS internalisation, with and without a 10 min preincubation, in PfATP2-HA knockdown and Control parasites.*** *NBD-PS internalisation was measured over 9 min at 15˚C in isolated trophozoite-stage parasites suspended in pH 7.1 Physiological Saline. The cells were either exposed to 5 mM GlcN for two days in the lead up to the experiment to reduce PfATP2-HA expression (+GlcN; pink) or not exposed to GlcN (-GlcN; blue). GlcN was not present during the measurements. The bars and error bars show the mean ± SEM (from three independent experiments performed on different days, with all conditions tested concurrently) and the symbols show the data from individual experiments. The data are expressed as a percentage of the NBD-PS fluorescence (geometric mean) measured in the -GlcN Control. To determine whether preincubation had a significant effect on the activity of any of the compounds, two-way ANOVAs were performed for each compound using the normalised data. Preincubation did not significantly affect the activity of vanadate ($P$ = 0.2), MMV665794 ($P$ = 0.09) or MMV007224 ($P$ = 0.3).*

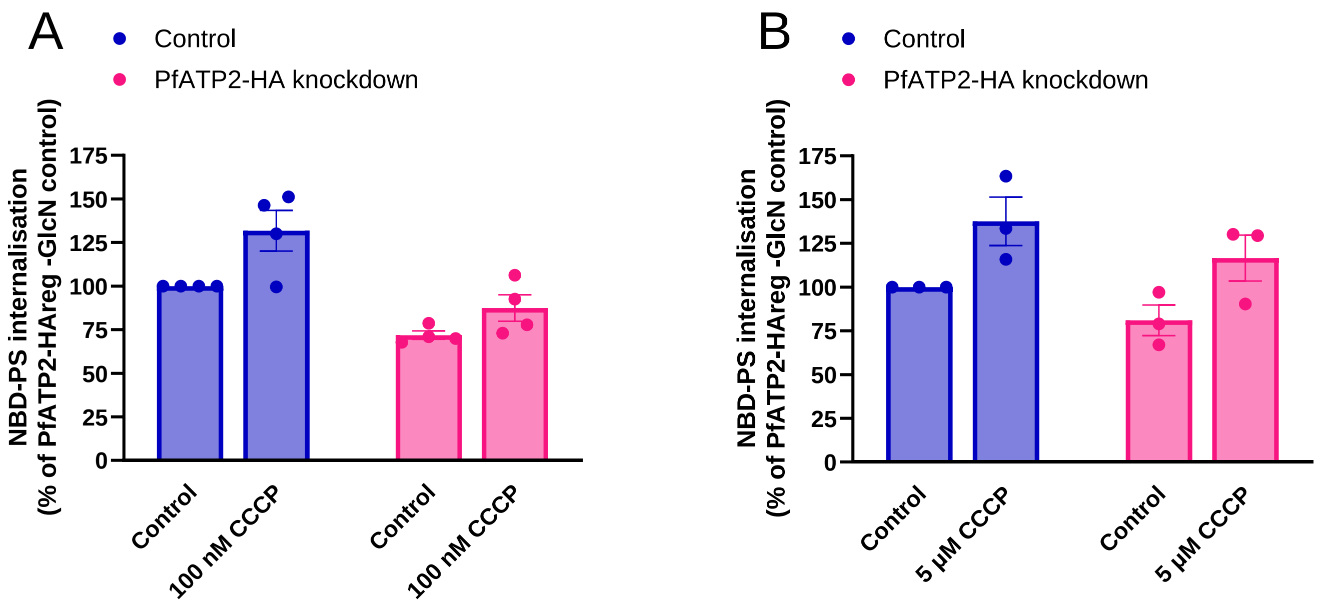


***Figure S8. CCCP did not inhibit the internalisation of NBD-PS by Control or PfATP2 knockdown parasites when tested at 100 nM (A) or 5 µM (B).*** *NBD-PS internalisation by isolated trophozoite-stage parasites was measured at 15˚C over 9 min. Symbols show data from each individual experiment with Control parasites (PfATP2-HAreg -GlcN; blue) and PfATP2-HA knockdown parasites (exposed to 5 mM GlcN for two days; pink). The bars and error bars show the mean ± SEM from four (**A**) or three (**B**) independent experiments (each performed on different days). The data are expressed as a percentage of the NBD-PS fluorescence (geometric mean) observed in Control parasites that were not treated with CCCP (0.1% v/v DMSO; solvent control). Two-way ANOVAs were performed with the natural logarithm transformed pre-normalised data. The P values for the effects of 100 nM CCCP and 5 µM CCCP were 0.07 and 0.03, respectively. In two experiments, 5 µM CCCP was tested alongside other compounds for which data are shown in **Fig. S7**. Thus, the Control data for two of the experiments in **Fig. S8B** are the same as those shown in **Fig. S7**.*

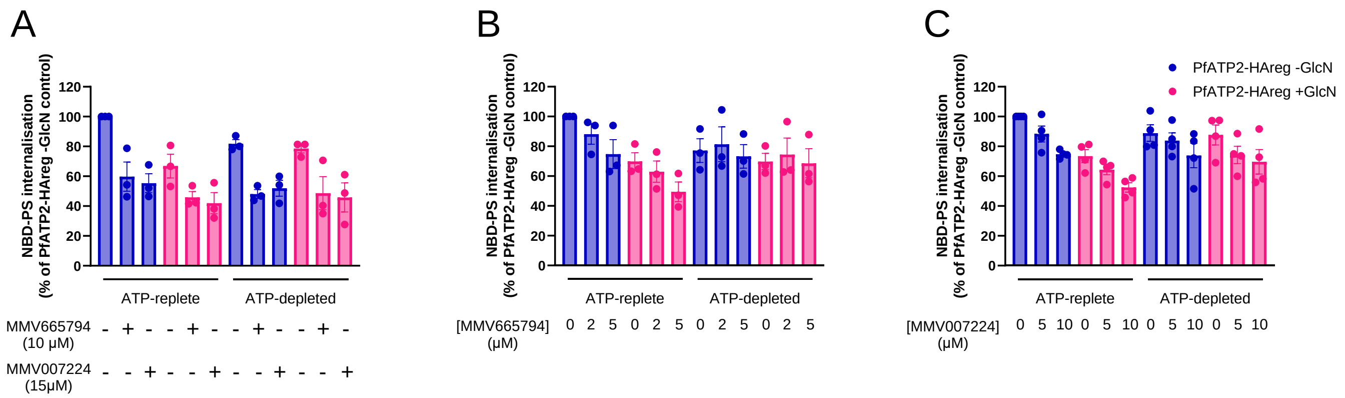


***Fig. S9: MMV665794 and MMV007224 inhibit NBD-PS internalisation by ATP-replete and ATP-depleted parasites with different potencies.*** *NBD-PS internalisation was measured over 9 min in isolated trophozoite-stage parasites suspended in pH 7.1 Physiological Saline (ATP-replete) or Glucose-free Saline (ATP-depleted) at 15˚C, in the absence (0.1% v/v DMSO; solvent control) or presence of the compounds and concentrations indicated. PfATP2-HAreg parasites were either exposed to 5 mM GlcN for two days in the lead up to the experiment to reduce PfATP2-HA expression (+GlcN; pink) or not exposed to GlcN (-GlcN; blue). GlcN was not present when parasites were exposed to NBD-PS. The data are expressed as a percentage of the NBD-PS fluorescence (geometric mean) measured in the ATP-replete parasites that were not exposed to GlcN or test compound. The data are from three (**A,B**) or four (**C**) independent experiments, in which all the conditions/parasite lines shown within one panel were tested concurrently. The symbols show the data from individual experiments; the bars and error bars show the mean ± SEM. The solvent control data in **A** are the same as three of those shown in **Fig. 5D** (as vanadate and the MMV compounds were tested together in three experiments). The data shown in the **Fig. 6** insets were derived from the same experiments as the data shown in this Figure (but in the case of the **Fig. 6** insets were normalised to the data for ATP-depleted parasites that were not exposed to GlcN or test compound).*

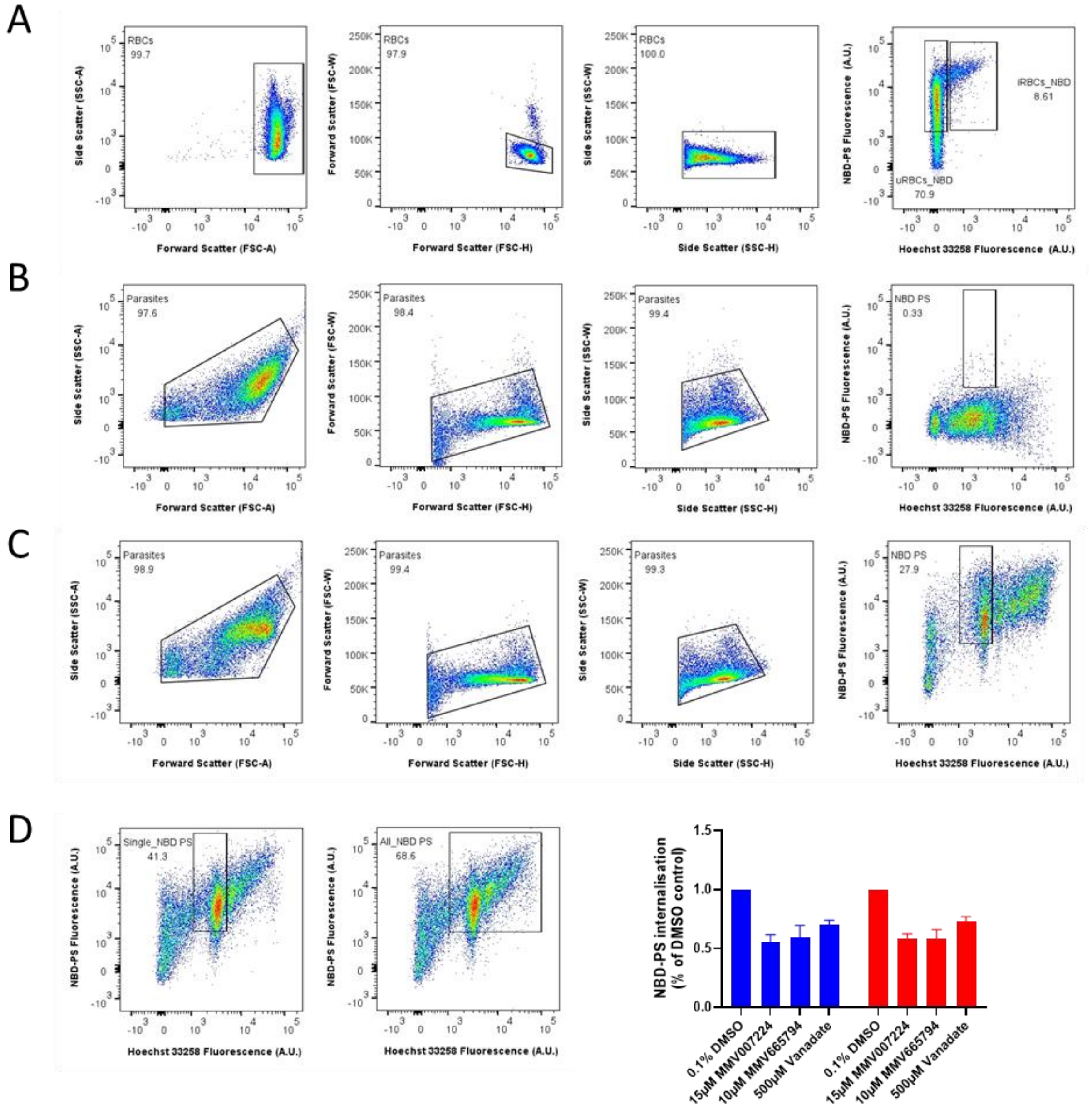


***Fig. S10 Gating strategy used to quantify the internalisation of NBD-PS by flow cytometry.*** *RBCs (**A**; uninfected and infected with* P. falciparum*) or isolated parasites (PfATP2-HAreg -GlcN; **B**-**D**) were first gated in a plot of SSC-A versus FSC-A to exclude debris. Cells were then gated in plots of FSC-W versus FSC-H and SSC-W versus SSC-H, in order to exclude doublets and aggregates. The cells were then gated in a plot of Hoechst 33258 fluorescence and NBD-PS fluorescence. The data in each panel are from a single experiment, representative of three or more independent experiments. **(A)** Gating strategy for uRBCs and iRBCs (infected with PfATP2-HAreg parasites) (-GlcN, 0.1% v/v DMSO condition). (**B-C**) Gating strategy for isolated trophozoite-stage parasites (PfATP2-HAreg -GlcN) that were not exposed to NBD-PS (**B**) or that were exposed to NBD-PS for 9 min at 15˚ (**C**). The gating strategy shown in panel **C** was used in this study. However, forming a larger gate encompassing parasites with varying Hoechst 33258 fluorescence levels (which are expected to be more diverse in level of maturity, with some parasites likely having multiple nuclei) yielded similar results (**D**). In the bar graph, the results obtained using the smaller gate shown on the left (‘Single_NBD PS’) are shown in blue, and those obtained using the larger gate shown on the right (‘All_NBD PS’) are shown in red. The data are for ATP-replete PfATP2-HAreg Control parasites (-GlcN) that were exposed to DMSO (0.1%; solvent control), 15 μM MMV007224, 10 μM MMV665794 or 500 μM vanadate (data from the same experiments are also shown in **Fig. 5D** and **Fig. S9A**). The bars and error bars show the mean + SEM from three independent experiments.*
